# OptoLoop: An optogenetic tool to probe the functional role of genome organization

**DOI:** 10.1101/2025.11.06.686574

**Authors:** Martin Stortz, Adib Keikhosravi, Gianluca Pegoraro, Tom Misteli

## Abstract

The genome folds inside the cell nucleus into hierarchical architectural features, such as chromatin loops and domains. If and how this genome organization influences the regulation of gene expression remains only partially understood. The structure-function relationship of genomes has traditionally been probed by population-wide measurements after mutation of critical DNA elements or by perturbation of chromatin-associated proteins. To circumvent possible pleiotropic effects of such approaches, we have developed OptoLoop, an optogenetic system that allows direct manipulation of chromatin contacts by light in a controlled fashion. OptoLoop is based on the fusion between a nuclease-dead SpCas9 protein and the light-inducible oligomerizing protein CRY2. We demonstrate that OptoLoop can drive the induction of contacts between genomically distant, repetitive DNA loci. As a proof-of-principle application of OptoLoop, we probed the functional role of DNA looping in the regulation of the human telomerase gene *TERT* by long-range contacts with the telomere. By analyzing the extent of chromatin looping and nascent RNA production at individual alleles, we find evidence for looping-mediated repression of *TERT*. In sum, OptoLoop represents a novel means for the interrogation of structure-function relationships in the genome at single-allele resolution.

## INTRODUCTION

The meter-scaled human genome needs to be tightly packaged inside the micrometer-scaled cell nucleus while allowing the correct execution of a variety of genome-associated functions, including gene expression. The highly complex organization of the genome at different length scales gives rise to distinct architectural features from nucleosomes and chromatin loops at the local level of organization to the formation of chromatin domains and chromosome territories for global organization (Misteli, 2020). These structural features have been widely implicated in genome function and gene regulation, including in cellular differentiation, development, and disease (Hnisz et al., 2016; Kragesteen et al., 2018; Oudelaar et al., 2020; Winick-Ng et al., 2021).

The organizational features of the 3D genome arise from an extensive network of chromatin-chromatin interactions across the chromatin polymer (Parmar et al., 2019; Conte et al., 2023). These contacts and the precise folding of the genome are regulated by specific DNA sequences and architectural proteins such as the CCCTC-binding factor (CTCF) and the molecular motor complex cohesin (Rowley and Corces, 2018; Mach et al., 2022; Uyehara and Apostolou, 2023). As a result of chromatin-chromatin interactions, distal genomic regions are brought into close physical proximity, forming chromatin loops which can span from a few kilobases to many megabases (Grubert et al., 2020). Chromatin loops are thought to play a prominent role in the regulation of gene expression by promoting contacts between genes and distal regulatory elements (Fulco et al., 2019; Popay and Dixon, 2022; Yang and Hansen, 2024).

The precise role of chromatin structural features in gene regulation is, however, not fully understood. On the one hand, there are prominent examples supporting a prominent role of genome organization in genome function, such as the observation of coupled transcription of paralogous genes with shared enhancers in *Drosophila* embryos through long-range promoter-promoter loops (Levo et al., 2022). Similarly, the activation of the *Shh* developmental gene (Kane et al., 2022) and the expression of key genes in neuron maturation rely on cohesin-mediated long-range enhancer-promoter loops (Calderon et al., 2022). On the other hand, other observations suggest that the relevance of genome structure for regulation of gene transcription is only limited. For instance, acute depletion of cohesin or CTCF leads to widespread disruption of chromatin loops, but to only modest alterations in gene regulation (Rao et al., 2017; Nora et al., 2017). Along the same lines, dorsoventral patterning during *Drosophila* embryogenesis involves major changes in gene expression, but chromatin organization, including relevant enhancer-promoter loops, remain unchanged during this process (Ing-Simmons et al., 2021). Furthermore, single-allele live cell imaging does not show any association between bursting of the developmental gene *Sox2* and physical proximity with its essential enhancer in embryonic stem cells (Alexander et al., 2019).

The apparent discrepancies in these observations may originate from the experimental challenges involved in manipulating chromatin structure and probing its effects on gene expression. The structure-function relationship of genomes has traditionally been explored by correlative studies observing natural changes in genome structure through development, differentiation, evolution, and disease (Krefting et al., 2018; Oudelaar et al., 2020; Akdemir et al., 2020; Winick-Ng et al., 2021; Hsieh et al., 2022; Liu et al., 2023) or by perturbing architectural drivers of genome architecture such as mutation of DNA sequences or manipulation of chromatin protein levels (Lupianez et al., 2015; Narendra et al., 2015; Rao et al., 2017; Hyle et al., 2019; Chakraborty et al., 2023). However, these approaches are limited in that they have pleiotropic effects that may affect gene activation independently from genome structure changes (Horsfield, 2023; Corin et al., 2025). Furthermore, these experiments usually rely on results obtained from biochemical, bulk-population measurements, lacking the precision of single-cell or single-allele level data. Finally, the perturbations used often occur at long time scales leading to possible secondary effects (Lupianez et al., 2015; Narendra et al., 2015; Chakraborty et al., 2023).

In recent years, attempts have been made to develop methods that overcome these limitations by synthetic manipulation of genome structure without mutating key architectural proteins or sequences (Deng et al., 2012; Morgan et al., 2017; Kim et al., 2019; Qin et al., 2022; Du et al., 2022). These methods are generally based on the dimerization of chromatin-binding proteins, such as zinc-finger proteins or bacterial Lac/Tet repressor proteins, directed to select target sequences. However, these approaches are often not inducible at short time scales, have not been shown to robustly induce long-range loops, nor do they generally provide single-cell or single-allele level information. To overcome these limitations, we have developed OptoLoop, an optogenetic method based on CRISPR/dCas9 tethering and light-dependent CRY2 oligomeric clustering, to rapidly induce chromatin loops with light. By pairing OptoLoop with a high-throughput imaging readout, we obtained single-cell and single-allele information to demonstrate the ability of this technique to induce looping between distant (> 1 Mb), repetitive, endogenous DNA loci at short time scales (> 90 min). We also provide proof-of-principle evidence that OptoLoop is a useful experimental platform to study the functional consequences of manipulating a biologically relevant chromatin loop in a controlled fashion using the human *TERT* gene as a model system.

## RESULTS

### Optimization of light-controlled clustering of CRY2

We sought to develop an optogenetic approach to rapidly induce chromatin looping in a controlled fashion in living human cells. For this purpose, we leveraged the light-controlled clustering properties of CRY2, a protein from *Arabidopsis thaliana* that forms oligomers and higher-order clusters in cells within minutes in response to pulses of blue light in a reversible manner (Fig. 1A) (Mas et al., 2000; Ozkan-Dagliyan et al., 2013; Ma et al., 2020; Palayam et al., 2021).

**Fig. 1.**
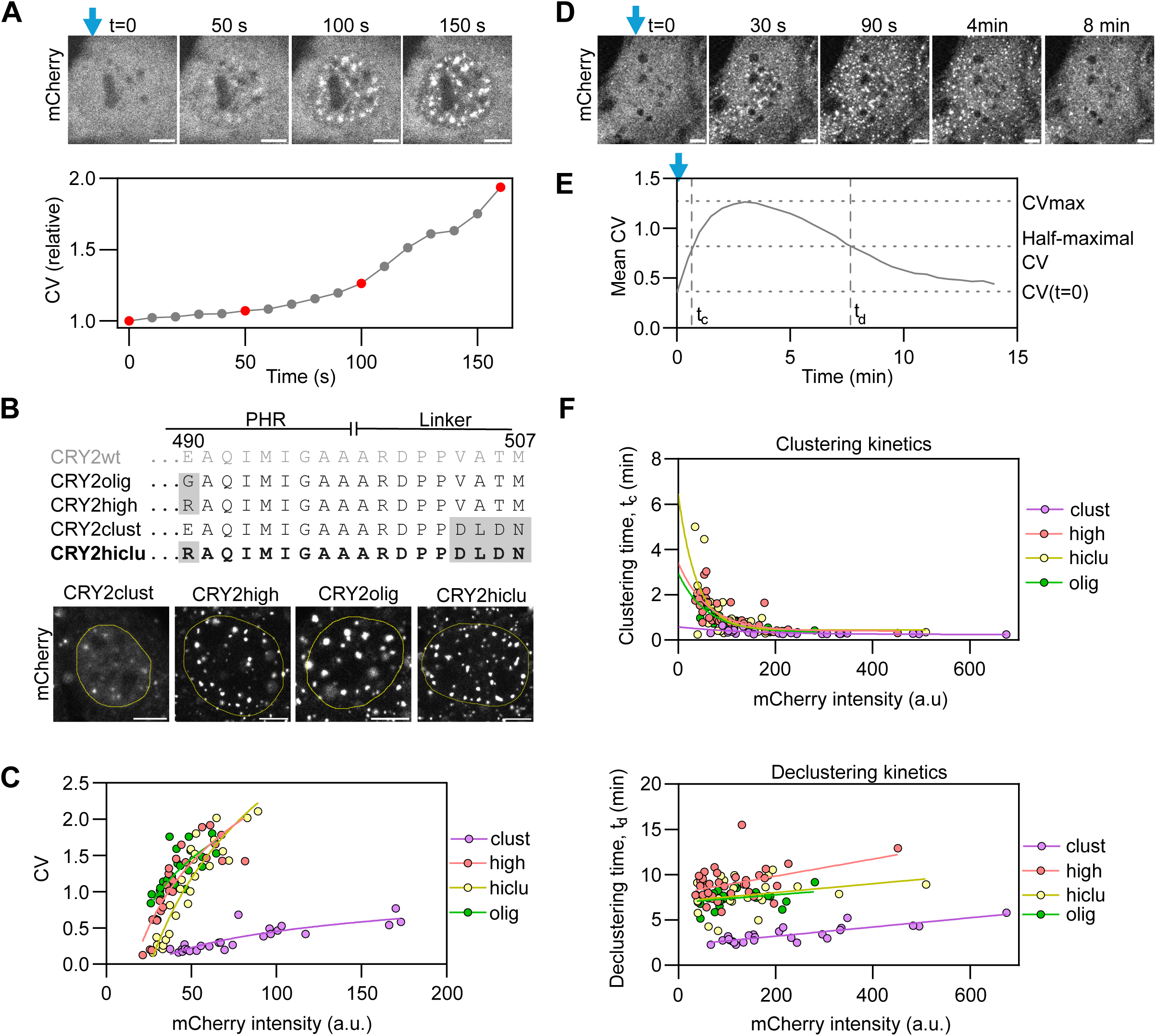
Exploration of optogenetic clustering properties of CRY2. **A)** Top panels: Time-lapse images of a NIH3T3 cell expressing CRY2high-mCherry activated with a 488-nm microscope laser starting at time t=0 (Blue vertical arrow, 1.5 s pulses every 10 s). Scale bar: 5 µm. Bottom panel: Coefficient of variation (CV) of fluorescence intensity calculated as the ratio between the nuclear intensity standard deviation and the nuclear intensity mean, presented as relative to the CV at time t=0. Data points corresponding to the images are marked in red. **B)** Top panel: Protein sequence of the C-terminus of the CRY2 PHR domain and part of the artificial linker used for C-terminal fusions for wild-type CRY2 (CRY2wt) and for CRY2 mutants. The newly generated variant CRY2hiclu is marked in bold. Mutations relative to the CRY2wt sequence are highlighted in gray. Bottom panel: Images of NIH3/T3 cells expressing CRY2 mutants fused to mCherry, illuminated with 1 s blue light pulses every 10 s for 15 min, and then fixed. The nucleus is delimited with a yellow line. Scale bar: 5 µm. **C)** CV calculated from images obtained from NIH3T3 cells expressing CRY2 variants fused to mCherry, illuminated with pulsed blue light for 15 min, and then fixed, plotted as a function of mCherry nuclear intensity. ∼25 cells were analyzed per sample (each dot represents one cell). Continuous lines represent simple logistic fits. **D)** Time-lapse images of a NIH3T3 cell expressing CRY2hiclu-mCherry activated once with the 488-nm microscope laser for 15 s at time t=0 (marked with a blue arrow). Scale bar: 5 µm. **E)** Mean CV calculated from time-lapse images obtained from NIH3T3 cells expressing CRY2olig-mCherry, illuminated with blue light at time t=0, and then kept without blue light. The clustering (t_c_) and declustering times (t_d_) were determined from individual kinetic curves. **F)** t_c_ (top panel) and t_d_ (bottom panel) represented as a function of mCherry nuclear intensity. ∼25-40 cells were analyzed per sample (each dot represents one cell). Continuous lines represent simple exponential (clustering) and linear (declustering) fits.

The wild-type CRY2 protein (CRY2wt) contains a photolyase homology region (PHR) domain which is sufficient for the light-inducible clustering of CRY2 (Taslimi et al., 2014). However, since the wild-type PHR domain has only poor clustering properties (Park et al., 2017), we tested previously described engineered mutants of CRY2 that demonstrate enhanced clustering capabilities upon light activation (Fig. 1B) (Taslimi et al., 2014; Park et al., 2017; Duan et al., 2017). All these variants have modifications in the PHR C-terminus: CRY2olig (Taslimi et al., 2014) and CRY2high (Duan et al., 2017) include a single E490G or E490R point-mutation, respectively (Fig. 1B), whereas CRY2clust (Park et al., 2017) contains a C-terminal DLDN motif (Fig. 1B). In addition, we also generated and tested a new CRY2 mutant, termed ‘CRY2hiclu’, which combines the E490R point-mutation of CRY2high and the C-terminal DLDN motif of CRY2clust (Fig. 1B).

To identify an optimal CRY2 variant for our purposes with high light-sensitivity and enhanced clustering properties, we quantified the degree of clustering for all engineered mutants from confocal images by measuring the CV of CRY2-mCherry fusions (Fig. 1A; see ‘Materials and Methods’). As a quantitative measurement of CRY2-mCherry clustering, the coefficient of variation of fluorescence intensity (CV) was measured for each cell nucleus, calculated as the ratio between the standard deviation of the CRY2 variant intensity and the mean intensity of the entire cell nucleus (Fig. 1A; see ‘Materials and Methods’). Unlike spot counting, calculation of CV does not require arbitrary thresholds for segmentation, providing a more accurate overall proxy for clustering when analyzing populations of clusters in a wide range of sizes and brightnesses as is the case with CRY2 clusters. To this end, we imaged NIH3T3 cells transiently expressing CRY2olig, CRY2high, CRY2clust, or CRY2hiclu fused to mCherry (Fig. 1B). We analyzed the degree of clustering achieved by each variant after illuminating cells with repeated 1 s pulses of blue light for 15 min at 10 s intervals (Fig. 1C). As previously reported, the clustering properties of CRY2 are dependent on its concentration (Taslimi et al., 2014; Park et al., 2017). We observed that CRY2high, CRY2olig, and CRY2hiclu achieve similar clustering levels but higher clustering than CRY2clust, reaching CV values ∼4-5 times larger than CRY2clust at comparable intermediate/high expression levels (Fig. 1C).

Light-induced CRY2 clustering is reversible, and clusters spontaneously disassemble in the absence of blue light (Taslimi et al., 2014). We measured the kinetics of assembly and disassembly of clusters of the engineered CRY2-mCherry mutants by acquiring time-lapse images of live cells immediately after a one-time activation of CRY2 with the microscope laser for 15 s and for up to 7-10 min after the light pulse (Fig. 1D; see ‘Materials and Methods’). We quantitatively characterized CRY2 clustering kinetics by estimating a half-maximal clustering time (t_c_) and a half-minimal declustering time (t_d_) (Fig. 1E). Clustering kinetics were similar for all variants, except CRY2clust which showed slightly faster clustering than the other variants (Fig. 1F). For cluster disassembly, CRY2clust also exhibited faster declustering kinetics (t_d_ ≈ 3.4 min) than the other variants (t_d_ ≈ 7.4-9.0 min) (Fig. 1F). We conclude that CRY2high, CRY2olig, and CRY2hiclu behave similarly, whereas CRY2clust presents lower clustering ability and faster deactivation kinetics, resulting in overall fewer steady-state clusters. For the purposes of generating a chromatin-looping tool, we used CRY2hiclu (hereafter ‘CRY2’), because it provides high and stable clustering properties at low concentrations.

### Manipulation of chromatin looping by light

We designed ‘OptoLoop’, an optogenetic tool to manipulate chromatin contacts controlled by light. OptoLoop is based on light-induced oligomerization of CRY2 fused to dCas9, a catalytically inactive version of the SpCas9 endonuclease, which can be tethered to target genomic locations via specific single-guide RNAs (sgRNAs) (Fig. 2A). In this approach, dCas9 is used to selectively target two genome regions of interest via use of sequence-specific sgRNAs, while the CRY2 moiety of the fusion protein brings together and stabilizes contacts between these chromatin regions by light-controlled CRY2-CRY2 oligomerization (Fig. 2A). We reasoned that use of a multivalent, oligomerizing protein such as CRY2 would be more efficient than previously used dimerizing proteins (Deng et al., 2012; Morgan et al., 2017; Qin et al., 2022; Du et al., 2022).

**Fig. 2.**
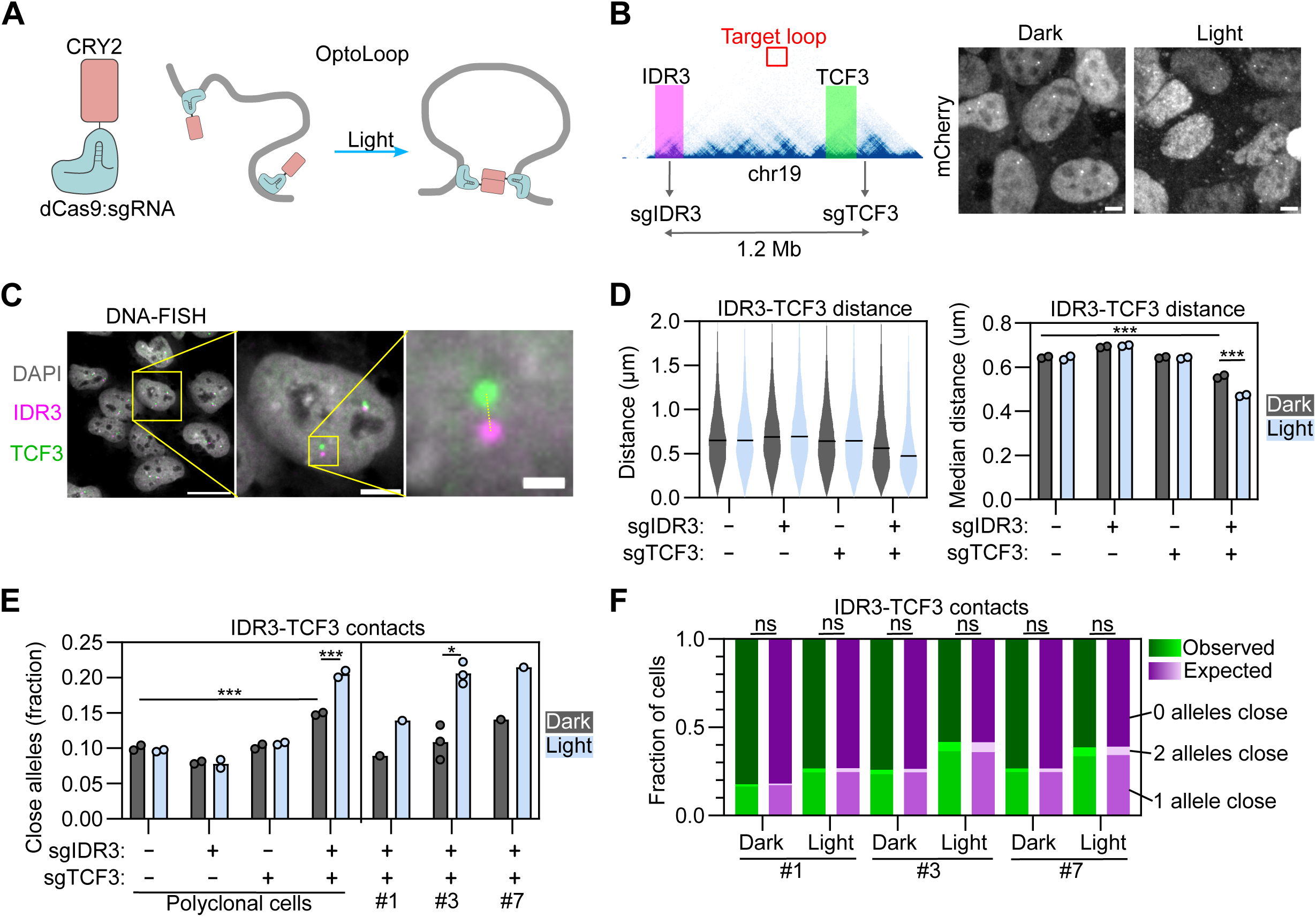
Optogenetic manipulation of contacts between repetitive genomic loci. **A)** Scheme of OptoLoop consisting of a fusion between dCas9 and the optogenetic protein CRY2. OptoLoop is targeted to specific genomic loci by introducing specific sgRNAs. CRY2-CRY2 interactions activated by blue light bridge targeted loci to form a chromatin loop. **B)** Left panel: Region of chromosome 19 showing sgIDR3 and sgTCF3 target sites, representative Hi-C contact map (data from (Rao et al., 2014)), and BACs used in DNA-FISH to label the *IDR3* (magenta) and *TCF3* loci (green). Right panel: mCherry-channel images of U2OS-dCas9-3XmCherry-CRY2 cells transfected with sgIDR3 and sgTCF3, kept in dark or illuminated with blue light for 3 h (1 s pulses every 10 s), and fixed. Scale bar: 5 µm. **C)** Left panel: Representative image of DNA-FISH for *IDR3* and *TCF3* with specific BAC FISH probes in U2OS cells. Right panel represents a single cell highlighted in left panel (yellow box), inset shows a single allele in this cell. Dashed line denotes the distance between the two FISH spots. Scale bars: 20 µm (left panel), 5 µm (right panel), 1 µm (inset). **D)** *IDR3-TCF3* distances, calculated for U2OS-dCas9-mCherry-CRY2 polyclonal cells transfected with indicated combinations of sgIDR3 and sgTCF3, kept under dark or illuminated for 3 h (1 s pulses every 10 s). Violin plot corresponds to a representative experiment, with black lines representing median distances. Bar plot represents means of two independent experiments. Each dot represents the median of typically 5000-10000 alleles analyzed per experiment. **E)** Fraction of alleles with *IDR3-TCF3* distance < 0.27 µm measured from DNA-FISH images for U2OS-dCas9-mCherry-CRY2 polyclonal cells and three clones of U2OS-dCas9-3XmCherry-CRY2 cells, transfected with indicated combinations of sgIDR3 and sgTCF3, and kept in dark or illuminated for 3 h (1 s pulses every 10 s). Each dot represents the fraction of typically 5000-10000 alleles analyzed per experiment. Bars represent means of two independent experiments. **F)** Measurement of cell-to-cell heterogeneity in loop formation. Bars with green shades: Observed fraction of cells with none, one, or both alleles with *IDR3-TCF3* distance < 0.27 µm obtained from a representative experiment shown in (E) with 2500-5000 cells analyzed per sample. Bars with magenta shades: Expected fraction of cells with none, one, or both alleles with *IDR3-TCF3* distance < 0.27 µm assuming that alleles from a same cell are independent between each other (Eqn. 2). Asterisks indicate significantly different comparisons (* for p<0.05, ** for p<0.01, *** for p<0.001).

We hypothesized that OptoLoop would require multiple tethering sites in each of the target genomic regions to sufficiently anchor the dCas9-CRY2 fusion protein to DNA and form stable chromatin contacts. Therefore, we decided to target endogenous tandem DNA repeats allowing us to use one unique sgRNA to tether the dCas9 module to tens of sites in a relatively small genomic region (Fig. 2B). For this purpose, we selected a pair of repetitive human loci, *IDR3* and *TCF3,* that can be efficiently labeled with a dCas9-based system as previously shown (Fig. S1A) (Ma et al., 2018). Each of these loci can be targeted with a unique sgRNA which binds to > 30 sites within a ∼2 kb region (Fig. 2B). *IDR3* and *TCF3* are separated by 1.2 Mb on chromosome 19 and they do not interact according to Hi-C contact maps (Fig. 2B, Hi-C data extracted from (Rao et al., 2014)). Accordingly, they were observed as spatially distant loci in the nucleus when imaged by dCas9-based labeling with an average separation distance of ∼1 µm (Ma et al., 2018). Based on these properties, we selected them as an ideal pair of loci to test the ability of OptoLoop to induce de novo long-range chromatin contacts.

To monitor our ability to induce chromatin loops at the single-allele level, we used a microscopy-based approach to measure *IDR3-TCF3* physical distances. Polyclonal U2OS cells stably expressing dCas9-mCherry-CRY2 were transfected with synthetic sgRNAs targeting *IDR3* and *TCF3* at high efficiency (Fig. S1B), and cells were illuminated with blue light pulses of 1 s every 10 s over a period of 3 h to induce looping (Fig. 2B). The positions of the two loci were determined by DNA-FISH using specific BACs (Fig. 2B-C) followed by high-throughput imaging and measurement of center-to-center distance between DNA-FISH signals for *IDR3* and *TCF3* at individual alleles (Fig. 2C; see ‘Materials and Methods’). We analyzed both the median *IDR3-TCF3* distance (Fig. 2D) and the fraction of alleles with *IDR3-TCF3* contacts defined as those below a distance threshold of 0.27 µm, similar to previously used thresholds to define chromatin interactions by confocal imaging (Fig. 2E; see ‘Materials and Methods) (Finn et al., 2019). We observed that light-activation decreased the median *IDR3-TCF3* distance from 0.56 to 0.47 µm (p-value = 0.00002; Fig. 2D) and increased the fraction of *IDR3-TCF3* contacts from 15% to 21% (p-value = 0.00003; Fig. 2E) only when both sgRNAs were introduced. The presence of each sgRNA alone did not induce *IDR3-TCF3* proximity (Fig. 2E). Extending the light-treatment from 3 h to 6 h did not increase the number of *IDR3-TCF3* contacts (Fig. S1C). A slight baseline OptoLoop activity independent of light was noted, suggesting that the presence of the two sgRNAs alone has a mild looping effect (Fig. 2D-E). These data suggest that OptoLoop can be used as an optogenetic tool to induce the formation of long-range chromatin contacts between spatially distant loci.

We hypothesized that OptoLoop expression levels would be relevant to its performance as oligomerization is sensitive to CRY2 concentration (Taslimi et al., 2014). Since the experiments described above were done with a polyclonal cell line with variable dCas9-mCherry-CRY2 expression levels between cells, we isolated and selected three clones with low (clone #1), intermediate (clone #3), and high (clone #7) expression levels of dCas9-3XmCherry-CRY2 (Fig. S1D) and compared their performances by studying their ability to induce *IDR3-TCF3* contacts (Fig. 2E). Higher expression levels indeed increased both the baseline OptoLoop activity independent of light and increased the light-induced *IDR3-TCF3* contacts (Fig. 2E). We selected clone 3 with intermediate expression levels for further analysis (Fig. 2E).

We next asked whether the two alleles in the same cell nucleus respond in a similar fashion to OptoLoop-induced chromatin looping or showed independent behavior. To do so, we calculated the fraction of cells with none, one, or both *IDR3-TCF3* alleles in contact upon light-induced dCas9-CRY2 clustering in the presence of sgIDR3 and sgTCF3 (Fig. 2F). The distributions of cells for the analyzed conditions were not statistically different than the expected distributions assuming complete independence between the alleles in the same cell (Fig. 2F). This observation suggests that *IDR3-TCF3* distance heterogeneity in light-induced conditions is not determined by extrinsic factors (dCas9-CRY2 levels, cell cycle stage, sgRNA transfection, etc.) but by intrinsic, allele-dependent factors such as chromatin state or epigenetic modifications. This observation is in line with prior observations of allele-independence of native chromatin contacts (Finn et al., 2019).

Taken together, these results establish OptoLoop as an optogenetic method to manipulate chromatin looping between repetitive genomic loci.

### Benchmarking OptoLoop

Light-activated dynamic looping (LADL) is a previously developed optogenetic tool based on the light-induced oligomerization of CRY2 and its heterodimerization with the partner protein CIBN (Fig. 3A) (Kim et al., 2019). LADL was shown to induce a ∼0.5-Mb chromatin loop between the *Zfp462* gene and an enhancer in mouse embryonic stem cells as assessed by the population-based chromosome conformation capture carbon copy (5C) technique (Kim et al., 2019). To benchmark OptoLoop against LADL, we applied both methods to the *IDR3* and *TCF3* targets and assessed their looping performance at the single-allele level by analyzing DNA-FISH distances.

**Fig. 3.**
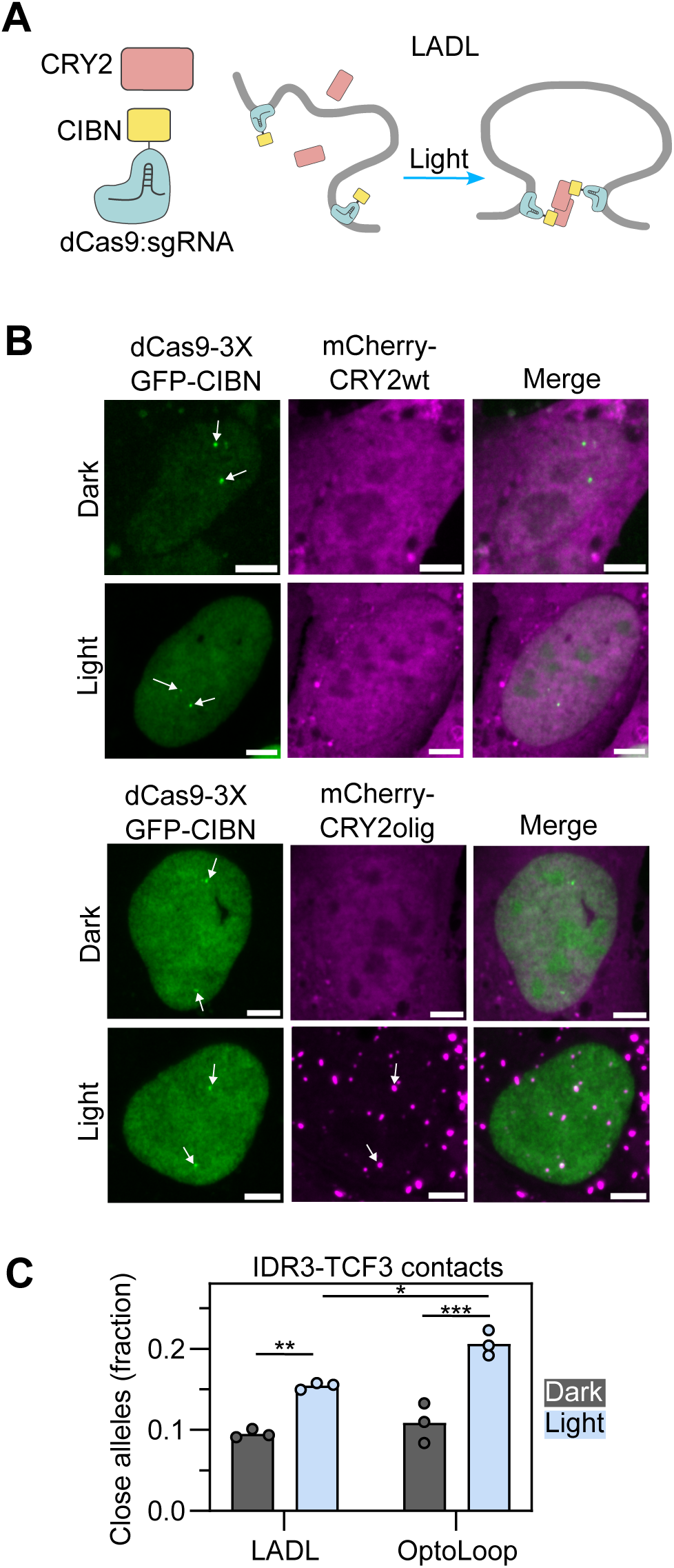
Benchmarking OptoLoop against a previous optogenetic manipulation tool. **A)** Scheme of LADL consisting of soluble CRY2wt and a fusion between dCas9 and the CRY2 partner CIBN. dCas9-CIBN tethers specific genomic loci and light-activation induces both CRY2-CRY2 and CRY2-CIBN interactions bridging the targeted loci to form a loop. **B)** Images of U2OS cells expressing dCas9-3XGFP-CIBN and mCherry fused to CRY2wt or CRY2olig, transfected with sgIDR3 and sgTCF3, and fixed after being kept under dark or illuminated with blue light pulses for 3 h (1 s pulses every 10 s). White arrows indicate spots corresponding to *IDR3/TCF3* loci. Scale bar: 5 µm. **C)** Fraction of alleles with *IDR3-TCF3* distance < 0.27 µm measured from DNA-FISH images of U2OS cell lines stably expressing dCas9-3XGFP-CRY2 and mCherry-CRY2olig (LADL, clone #7) or dCas9-3XmCherry-CRY2 (OptoLoop, clone #3), transfected with sgIDR3 and sgTCF3, and kept under dark or illuminated with blue light for 3 h (1 s pulses every 10 s). Each dot represents the fraction of 5000-7000 alleles analyzed per experiment. Bars represent the means of three independent experiments. Asterisks indicate significantly different comparisons (* for p<0.05, ** for p<0.01, *** for p<0.001).

For this purpose, we generated U2OS cell lines stably expressing both components of the LADL system fused to fluorescent proteins (dCas9-3XGFP-CIBN and mCherry-CRY2) to easily monitor their spatial distributions and expression levels (Fig. 3B). We tested LADL performance using wild-type CRY2 as originally described (Kim et al., 2019) and the enhanced clustering mutant CRY2olig. We verified labeling of the *IDR3 and TCF3* loci by dCas9-3XGFP-CIBN and light-induced recruitment of mCherry-CRY2olig to these locations (Fig. 3B). However, we did not detect recruitment of CRY2wt to target sites by imaging (Fig. 3B). Using polyclonal cell lines expressing varying levels of dCas9-3XGFP-CRY2 and mCherry-CRY2wt or –CRY2olig, we found that LADL performed better with CRY2olig than with CRY2wt (Fig. S2A) and that looping performance inversely correlated with CRY2 expression levels (Fig. S2A) which we also consistently observed across different single clones (Fig. S2B-C). Comparison of LADL and OptoLoop clones indicated that OptoLoop performed slightly, but significantly, better than LADL in bringing *IDR3* into proximity of *TCF3* (21% contacts vs 15% contacts, p-value = 0.012; Fig. 3C). We conclude that OptoLoop is an effective tool to induce chromatin looping between specific genomic loci.

### Use of OptoLoop to assess the biological function of chromatin looping

Next, we applied OptoLoop to probe the functional role of chromatin looping in the context of endogenous gene regulation. As a model system, we used the gene regulatory mechanism of “telomere position effect over long distance” (TPE-OLD) (Fig. 4A). In TPE-OLD, telomeres repress the expression of some genes located several Mbs away through long-range chromatin loops (Robin et al., 2014; Kim et al., 2016; Jager et al., 2022; Chevalier et al., 2025). Importantly, the repressive ability of telomeres on these distant genes is directly proportional to telomeres length, making this mechanism relevant for aging and cancer, when they can undergo shortening (Robin et al., 2014; Kim et al., 2016). One of the target genes of TPE-OLD is the human telomerase gene *TERT*, which is located ∼1.3 Mb away from the telomere on chromosome 5 (Fig. 4B) (Kim et al., 2016). In the TPE-OLD model, *TERT* is repressed via looping of the telomere to associate with *TERT* (Fig. 4A). Evidence of this mechanism comes from bulk-population measurements in which a decrease in telomere length correlated with a decrease in telomere:*TERT* contacts and up-regulation of *TERT* (Kim et al., 2016). Since this genomic region contains multiple endogenous tandem repeats, we wondered if we could use OptoLoop to manipulate the interactions of the *TERT* locus with its chromosome 5 p-arm telomere (Fig. 4B).

**Fig. 4.**
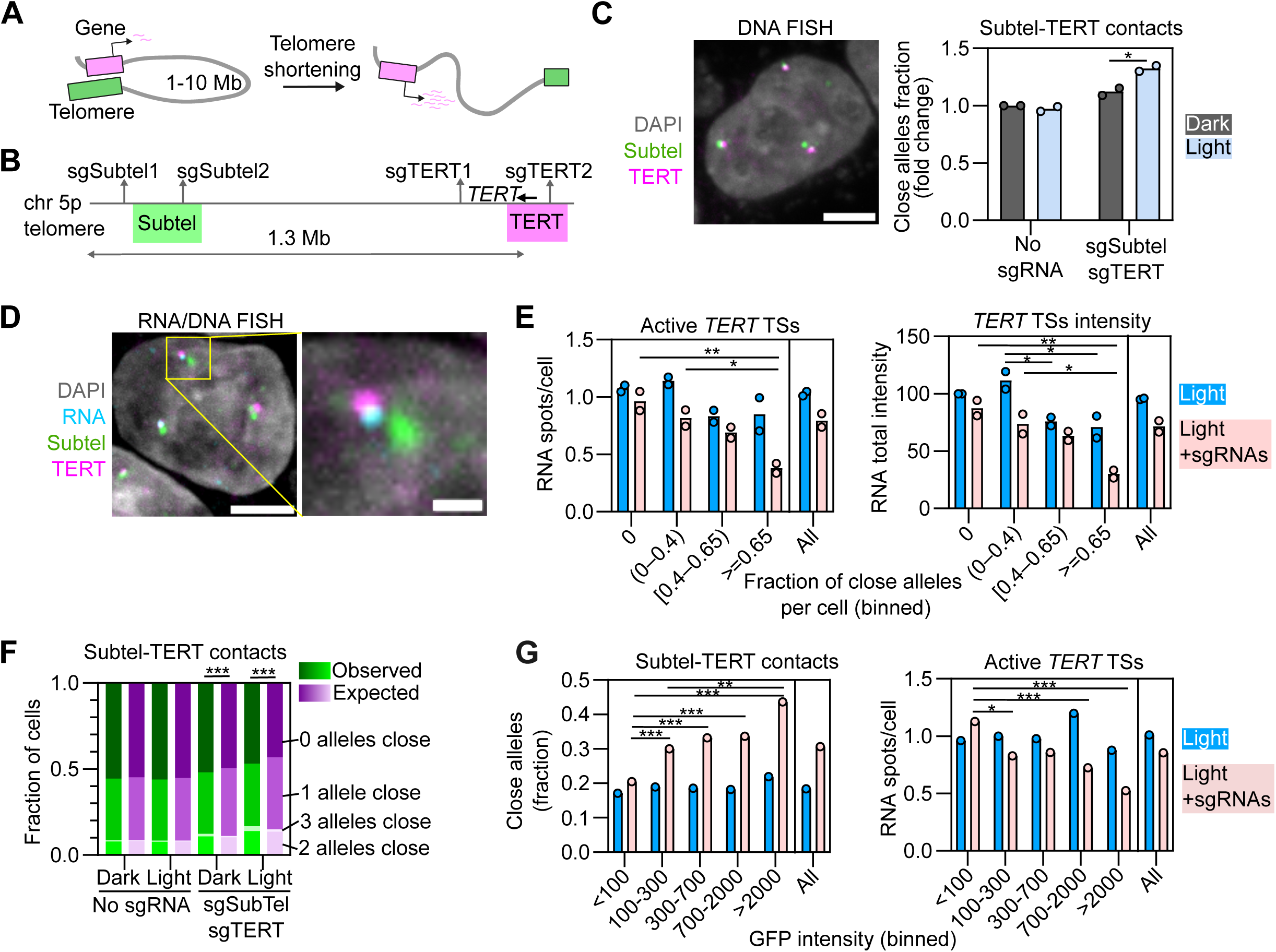
Functional effects of genome organization manipulation on gene regulation. **A)** Scheme of ‘telomere position effect over long distances’ model. Gene repression by long-range looping with telomere depends on telomere length. **B)** Region of chromosome 5 showing sgSubtel (sgSubtel1 + sgSubtel2) and sgTERT (sgTERT1 + sgTERT2) target sites, and BACs used in DNA-FISH to label the 5p subtelomeric region (Subtel, green) and the *TERT* locus (magenta). **C)** Left panel: Representative image of DNA-FISH with Subtel and *TERT* BAC probes in HeLa cells. Scale bar: 5 µm. Right panel: Fraction of alleles with Subtel-*TERT* distance < 0.27 µm measured from DNA-FISH images of HeLa cells stably expressing dCas9-3XGFP-CRY2, transfected with none or sgSubtel and sgTERT sgRNAs, and kept under dark or illuminated with blue light for 3 h (1 s pulses every 10 s). Each dot represents the fraction of typically 6000-20000 alleles analyzed per experiment. Bars represent the means of two experiments. Values are represented as relative to control (no sgRNAs, no light). **D)** Representative image of RNA/DNA-FISH with RNA probes against *TERT* pre-mRNA and Subtel and *TERT* BAC probes in HeLa. Inset shows a single allele (highlighted with a yellow square). Scale bars: 5 µm (left image), 1 µm (inset). **E)** Number of active *TERT* transcription sites (TSs) per cell (left panel) and total *TERT* pre-mRNA intensity (right panel) measured from RNA/DNA-FISH images from HeLa cells stably expressing dCas9-3XGFP-CRY2, transfected with none or sgSubtel and sgTERT sgRNAs, and illuminated with blue light for 4 h (1 s pulses every 10 s). Cells were binned according to their fraction of alleles with Subtel-*TERT* distance < 0.27 µm. Average measurements from the whole population of cells (i.e. not binned) are also shown. Each dot represents the mean of typically 150-1500 cells analyzed per bin, per experiment. Bars represent the means of two experiments. RNA total intensity was normalized to that of control (Bin ‘0’, no sgRNAs). **F)** Bars with green shades: Observed fraction of cells with none, one, two, or all three alleles with Subtel-*TERT* distance < 0.27 µm obtained from a representative DNA-FISH experiment shown in (C) with typically 6000-7000 cells analyzed per sample. Bars with magenta shades: Expected fraction of cells with none, one, two, or all three alleles with Subtel-*TERT* distance < 0.27 µm assuming that alleles from a same cell are independent of each other (Eqn. 3). **G)** Fraction of alleles with Subtel-*TERT* distance < 0.27 µm (left panel) and average number of active *TERT* transcription sites per cell (right panel) measured from a representative RNA/DNA-FISH experiment shown in (E). Cells were binned according to their mean GFP nuclear intensity. Average measurements from whole population of cells (i.e. not binned) are also shown. Each dot represents the fraction of 300-1800 alleles analyzed per bin (left panel) or the mean of 100-600 cells analyzed per bin (right panel). Asterisks indicate significantly different comparisons (* for p<0.05, ** for p<0.01, *** for p<0.001).

For this purpose, we targeted OptoLoop to two repetitive loci in the subtelomeric 5p region (sgSubtel) and two repetitive loci around the *TERT* gene (sgTERT) with the goal of bringing the *TERT* promoter into proximity to the 5p telomere in U2OS cells expressing dCas9-3XmCherry-CRY2 (Fig. 4B). Indeed, light-activation of OptoLoop induced *TERT*:subtelomere contacts (distance < 0.27 µm) in 50% of alleles compared to 36% in control cells kept in the dark (p-value = 0.006, Fig. S3B). Induction of contacts, albeit somewhat weaker, also occurred in HeLa cells (Fig. 4C, Fig. S3C-D), which have active telomerase and where *TERT* transcription is sensitive to the proposed TPE-OLD mechanism (Kim et al., 2016).

To probe for functional consequences of induced looping, we performed simultaneous RNA/DNA-FISH to measure *TERT* transcription at individual alleles after illuminating HeLa-OptoLoop cells with 1 s pulses at intervals of 10 s over 4 h (Fig. 4D, S4A-B). Based on preliminary RNA-FISH analysis, we compared light-treated cells transfected with none or with both sgSubtel and sgTERT (Fig. S4C). We performed single-cell analysis by calculating the fraction of alleles with *TERT*:subtelomere contacts (distance <0.27 µm) for every cell and binned cells accordingly (Fig. 4E). In the presence of sgRNAs targeting the loop anchors, cells with more DNA contacts also showed a 61% drop in the average number of *TERT* active transcription sites (p-value = 0.006 comparing cells with 0 vs ≥ 0.65 alleles close) and a 66% drop in the total *TERT* RNA signal intensity per cell (p-value = 0.003 comparing cells with 0 vs ≥ 0.65 alleles close) (Fig. 4E). In control cells without sgRNAs, a significant reduction in the total RNA signal intensity (p-value: 0.022) but not RNA spot number (p > 0.05) was observed (Fig. 4E). This data is consistent with *TERT* repression by *TERT*:telomere looping.

We noticed that the extent of the effect on RNA production varied between individual cells. To ask whether the heterogeneous response in *TERT* expression across the cell population was related to cell-to-cell heterogeneity in OptoLoop performance, we calculated the fraction of cells with zero, one, two, or three alleles with *TERT*:subtelomere contacts and compared them with the values expected for a completely independent behavior of alleles (Fig. 4F). We find that only cells transfected with sgRNAs, and especially those treated with light, had non-independent distributions (Fig. 4F). Specifically, OptoLoop activation led to more cells than expected with none or all three alleles close, and fewer cells than expected with one allele close compared to the random distribution (Fig. 4F). This result indicates that alleles from the same cell tend to respond more similarly to OptoLoop activation than alleles from different cells, partly explaining the cell-to-cell variability.

Finally, we hypothesized that cell-to-cell heterogeneity in OptoLoop performance and RNA production might be related to differences in dCas9-3XGFP-CRY2 expression levels within the cell population. Therefore, we measured GFP intensity simultaneously with RNA-FISH imaging and found that despite being a monoclonal cell line, expression levels were variable amongst individual cells in the population (Fig. S4D). As expected, DNA contacts and RNA production were not related to GFP intensity for cells not transfected with sgRNAs (Fig. 4G). However, for cells transfected with the sgRNAs, higher GFP intensities correlated with higher numbers of *TERT*:subtelomere contacts and with lower numbers of active *TERT* transcription sites (Fig. 4G). These results suggest that single-cell variability in OptoLoop efficiency is related to fluctuations in dCas9-CRY2 levels. These data also further support the association between the engineered loop and its repressive effect on *TERT* expression, demonstrating a functional effect of controlled manipulation of local genome organization.

## DISCUSSION

Here, we report the development of OptoLoop, an optogenetic tool to engineer synthetic chromatin loops and to interrogate the cause-effect relationship between 3D genome organization and gene regulation in mammalian cells. We describe its optimization, validate its ability to promote chromatin loops, and we provide proof-of-principle of its utility by demonstrating transcriptional regulation upon long-range loop formation at the human *TERT* locus, in support of previous reports of this regulatory mechanism (Robin et al., 2014; Kim et al., 2016).

At the center of OptoLoop is a fusion protein between a catalytically inactive CRISPR/dCas9 endonuclease which allows targeting of the fusion protein to select genomic sites via sgRNAs and the CRY2 protein which oligomerizes in response to blue light (Taslimi et al., 2014). OptoLoop is relatively simple in design and only requires the expression of the fusion protein, the transient transfection of synthetic sgRNAs targeting genomic loci of interest, and the stimulation with blue light to induce the formation of specific chromatin-chromatin contacts in a controlled fashion.

OptoLoop is highly versatile as the dCas9-CRY2 fusion protein can, in theory, be targeted to any genomic location using appropriate sgRNAs. Synthetic sgRNAs targeting loci of interest are commercially available and can be transfected into most cell lines at very high efficiency via lipofection. Since oligomerization properties are concentration-dependent dCas9-CRY2 levels and sgRNA transfection should be tested to maximize OptoLoop performance. Optogenetic activation of CRY2 with blue light can be achieved with a variety of setups and devices, including handheld lamps, microscope light sources or laser lines (Hernandez-Candia et al., 2019). In our studies, we observed best results in cell lines with stable expression of dCas9-CRY2 and in single-cell clones with homogeneous expression levels.

OptoLoop can be used alone to induce chromatin loops or can be combined with functional gene expression read-out assays such as FISH. We used DNA-FISH and nascent RNA-FISH to detect chromatin contacts and gene activity, respectively, followed by high-throughput imaging, routinely obtaining data from thousands of cells and alleles per sample. This strategy provides single-cell and single-allele level information, enabling probing the mechanistic relationship between observed changes in genome structure and function at individual alleles as well as in the population as a whole. As previously reported, chromatin contacts and gene bursting are highly variable across cell populations (Finn et al., 2019; Rodriguez et al., 2019). In line with cell-to-cell variability in 3D genome organization, we observed that induction of loops by light is incomplete, occurring in only a fraction of alleles, and that increasing the duration of the light treatment did not increase the number of contacts. In U2OS cells, we verified that the OptoLoop performance at the two alleles in the same cell was unrelated to each other. This means that each allele responds independently and that we cannot explain the incomplete performance of OptoLoop by single-cell properties such as cell cycle stage, dCas9-CRY2 expression, or sgRNA transfection. This finding is also in line with observations of allele-independence in chromatin looping (Finn et al., 2019). We can speculate that some alleles might be in an ‘incompetent’ state for looping, maybe related to chromatin structure or state. In contrast, we observed in HeLa cells that the behavior of alleles in the same nucleus was partially coordinated. Together with the observation that HeLa cells with higher dCas9-CRY2 levels had more alleles with synthetic loops, these data suggest that the inferior performance of OptoLoop in HeLa cells is at least partially explained by heterogeneous dCas9-CRY2 levels amongst cells in the population.

Several approaches to manipulate chromatin looping in eukaryotic cells have been reported (Deng et al., 2012; Morgan et al., 2017; Kim et al., 2019; Qin et al., 2022; Du et al., 2022). Use of these methods in follow-up studies has, however, been limited (Deng et al., 2014; Bartman et al., 2019; Benabdallah et al., 2019; Wang et al., 2021). Some of these methods were based on programmable DNA-binding proteins as targeting moieties and looping was not inducible (Deng et al., 2012; Benabdallah et al., 2019). Later, the use of CRISPR/dCas9 for tethering increased versatility (Morgan et al., 2017; Kim et al., 2019; Qin et al., 2022), and the development of inducible systems by either light (Kim et al., 2019) or a chemical compound (Morgan et al., 2017; Du et al., 2022) allowed more precise control of genome structure at shorter time scales. While most prior methods were based on dimerizing proteins, we hypothesized that use of an oligomerizing/clustering protein such as CRY2 would be more efficient at inducing chromatin loops. We benchmarked OptoLoop against its closest counterpart LADL, a previously reported looping tool also based on CRY2 optogenetic oligomerization (Kim et al., 2019). We extended the earlier bulk population-based biochemical studies on LADL by measuring its looping efficiency at the single-allele level. We verified a slightly superior performance of OptoLoop at the same locus. It is also worth noting that we have here used OptoLoop to induce longer-range loops (> 1 Mb) compared to the shorter loops of typically 10-100 kb previously manipulated in mammalian cells (Deng et al., 2012; Bartman et al., 2016; Morgan et al., 2017; Qin et al., 2022).

We have encountered some limitations of OptoLoop. First, as discussed above, we observed that not all alleles respond to OptoLoop activation by light. Although it is challenging to directly compare microscopy-based contact measurements with chromosome capture results, our single-allele analysis of LADL suggests that the increase in looping observed in population-based methods is driven by a subset of cells and does not reflect the looping behavior of the overall population. Second, we have demonstrated robust OptoLoop performance when targeting repetitive DNA loci, but our attempts to label nonrepetitive loci with dCas9 were not successful. This limitation is consistent with previous reports showing that targeting a tandem repeat of multiple sites with one sgRNA is more efficient than targeting the same number of unique sites with individual sgRNAs (Park and Kim, 2025). The reason behind the lower efficiency of targeting unique sites remains unclear but may be related to variable concentrations and local availability of each specific dCas9:sgRNA complex, possible competition between different dCas9:sgRNA species, and differences in sgRNA affinities to target sites. Third, we observed some OptoLoop baseline activity in the absence of light (i.e. increase in chromatin contacts by addition of sgRNAs only), and this baseline activity was more evident with higher dCas9-CRY2 levels. This observation emphasizes the need for optimization of dCas9-CRY2 expression levels to achieve high looping performance with minimal baseline activity. Also, proper precautions should be taken during experiments to avoid unwanted CRY2 activation in control conditions by keeping cells in the dark (Hernandez-Candia et al., 2019).

While we consider our study as a useful step towards creating synthetic biology tools for the study of chromatin architecture, future efforts are needed to improve synthetic looping tools and make them more robust, accessible and versatile. Recent innovations have made possible orthogonal, dual color dCas9-based labeling of nonrepetitive loci, allowing high temporal and spatial resolution (Ma et al., 2018; Clow et al., 2022; Lyu et al., 2022; Yang et al., 2024; Zhu et al., 2025). Integrating these dual color tethering systems with an optogenetic clustering module would allow assessing chromatin looping in dynamic experiments in individual cells in real time. Gene transcription could also be monitored in real time by coupling a compatible nascent RNA labeling system, such as the widely used MS2/PP7 RNA labeling approaches (Braselmann et al., 2020). Additionally, the capabilities of OptoLoop could be extended to the manipulation of multiway chromatin contacts based on the multivalent, nonstoichiometric nature of CRY2 clustering. Furthermore, OptoLoop could be combined with other tools to manipulate interactions with nuclear subcompartments, phase-separated condensates, and transcriptional machinery (Wang et al., 2018; Schneider et al., 2021; Kim et al., 2023; Strom et al., 2024). These synthetic biology approaches will be useful tools to answer the fundamental question of how genome organization contributes to genome function.

## MATERIALS AND METHODS

### Cell lines

NIH3T3 (mouse fibroblasts, ATCC #CRL-1658), U2OS (from human osteosarcoma, ATCC #HTB-96), HeLa (from human cervical adenocarcinoma, ATCC #CRM-CCL-2), and Lenti-X HEK-293T (from human embryonic kidney, cat. #632180 from Takara Bio, Japan) cell lines were cultured in Dulbecco’s modified Eagle’s medium (Gibco, Waltham, MA, USA) supplemented with 10% (15% for NIH3T3) fetal bovine serum (Gibco, Waltham, MA, USA) plus 100 IU/ml penicillin and 100 µg/ml streptomycin at 37°C in a humidified atmosphere with 5% CO_2_. Cell lines were routinely checked for Mycoplasma contamination.

### Plasmid constructs

A list of all plasmids with their corresponding sources can be found in Table S1. pCMV-CRY2hiclu-mCherry was obtained by replacing residues 504-507 from pCMV-CRY2high-mCherry by Asp-Leu-Asp-Asn via site-directed mutagenesis with the Q5 Site-Directed Mutagenesis kit (New England Biolabs, Ipswich, MA, USA). pHAGE-dCas9-mCherry-CRY2 was generated by replacing 2XmCherry in pHAGE-dCas9-3XmCh with PCR-amplified CRY2hiclu using NotI/XhoI sites. pHAGE-dCas9-3XmCherry-CRY2 and pHAGE-dCas9-3XGFP-CRY2 were generated with the NEB HiFi DNA assembly kit (New England Biolabs, Ipswich, MA, USA) by inserting PCR-amplified CRY2hiclu in XhoI-linearized pHAGE-dCas9-3XmCh and pHAGE-dCas9-3XGFP, respectively. pHAGE-dCas9-3XGFP-CIBN was generated by inserting PCR-amplified CIBN from pEF1a-dCas9-CIBN in pHAGE-dCas9-3XGFP using XhoI/Xba sites.

All plasmid constructs generated in this work were verified by whole-plasmid sequencing and will be deposited in Addgene.

### Generation of stable cell lines

Cell lines with stable expression of dCas9-3XGFP, dCas9-mCh-CRY2, dCas9-3XmCh-CRY2, dCas9-3XGFP-CRY2, dCas9-3XGFP-CIBN, mCh-CRY2wt, or mCh-CRY2olig were generated by lentiviral transduction. Lentiviral particles were generated by transfecting a subconfluent 10-cm cell culture dish of Lenti-X HEK-293T cells with 4 µg psPAX2, 1 µg pMD2.G, 4 µg of lentiviral construct, and 50 µL of 1 mg/mL polyethylenimine (Thermo Fisher, Waltham, MA, USA). After 48 h, supernatant with lentivirus was collected, filtered through a 0.45 µm filter, and concentrated overnight with Lenti-X Concentrator (Takara Bio, Shiga, Japan). Then, lentiviral particles were centrifuged (1500 xg, 45 min) and resuspended in growth media. Target cells were incubated with this lentiviral suspension and 10 µg/mL of Polybrene reagent (Sigma-Aldrich, Burlington, MA, USA) in subconfluent 6-well plates for 24 h.

Transduced cell lines were sorted for mCherry+ or GFP+ cells with a MA900 Cell Sorter (Sony, Tokyo, Japan) to obtain polyclonal cell lines with variable expression levels. Single-clones were isolated by seeding polyclonal cell lines at very low density in a 15-cm dish and transferring single-colonies with PYREX cloning cylinders (Corning, Corning, NY, USA) after 2 weeks. U2OS-dCas9-3XGFP and U2OS-dCas9-mCh-CRY2 cell lines were polyclonally sorted for low expression. U2OS-dCas9-3XGFP-CIBN/mCh-CRY2wt and U2OS-dCas9-3XGFP-CIBN/mCh-CRY2olig cell lines were polyclonally sorted for low GFP expression and into two populations with low and high mCherry expression. 12 clones were isolated from U2OS-dCas9-3XGFP-CIBN/mCh-CRY2olig (low mCherry population). U2OS-dCas9-3XmCherry-CRY2 cell line was sorted for mCherry positive cells and 9 clones were isolated. HeLa-dCas9-3XGFP-CRY2 cells were sorted into two populations with low and high GFP levels, and 7 clones were isolated. All data shown for HeLa cells was obtained from experiments done with one clone isolated from the high GFP population.

### Transient transfections

NIH3T3 cells were subjected to reverse-transfection with DNA plasmids by seeding 35,000 cells with 150 ng plasmid DNA and 0.75 µL Lipofectamine LTX (Thermo Fisher, Waltham, MA, USA) in Lab-Tek 8-well chambered coverglass (Thermo Fisher, Waltham, MA, USA). These cells were used for studying properties of different CRY2 variants.

Synthetic single-guide RNAs (sgRNAs) were custom-ordered from Synthego (Reedwood City, CA, USA) and are listed in Table S2. sgRNAs were synthesized with a 2’-O-Methyl modification at the 3 first and last bases, and 3’ phosphorothioate bonds between the 3^rd^ and 2^nd^ last bases for increased stability. RNA was introduced in the target cells by reverse-transfection. 9,000 U2OS cells or 16,000 HeLa cells were seeded with 1 or 0.5 pmol sgRNA (total) and 0.25 µL or 0.125 µL Lipofectamine 2000 (Thermo Fisher, Waltham, MA, USA), respectively, in a final volume of 50 µL growth media in a PhenoPlate 384-well microplate (Revvity, Waltham, MA, USA).

### Light activation

Except for studying CRY2 clustering/declustering kinetics, CRY2 activation was performed by illuminating cells from above the plate with a 455-nm LED source (M455L3-C4, ThorLabs, Newton, NJ, USA) operated by a LED Driver DC2200 (ThorLabs, Newton, NJ, USA). Illumination was performed at pulses of 1 s every 10 s. Power was set at 5% (for HeLa) or 10% (for NIH3T3 and U2OS) at a current of 1000 mA. The duration of the illumination treatments was variable for different experiments (see figure legends for details). For controls, wells with cells kept in the dark were covered with aluminum foil. Cells were fixed with 4% paraformaldehyde (Electron Microscopy Sciences, Hatfield, PA, USA) for 10 min at room temperature (RT) immediately after completion of the illumination protocol.

### Analysis of CRY2 variants properties

NIH3T3 cells cultured in 8-well coverglass and transfected with CRY2 variants fused to mCherry were kept in dark until the time of the experiment and imaged with a LSM880 confocal microscope (Zeiss, Oberkochen, Germany). For measuring fusion protein clustering, cells were illuminated with a 455-nm LED for 15 min and then fixed. The coefficient of variation (CV) was used as a measure of clustering and was calculated for individual cells as the ratio between the standard deviation of the intensity and the mean intensity of the entire cell nucleus.

For measuring clustering/declustering kinetics, live cells were illuminated at time t=0 by repeated scans with the 488-nm line of the Ar laser for 15 s, followed by time-lapse images every 30 s. CV was calculated for every time point. Clustering time (t_c_) for individual cells was estimated by linear interpolation as the time to reach half-maximal clustering as the difference between maximum CV and initial CV at time t=0. Declustering time (t_d_) for individual cells was estimated by linear interpolation as the time to reach again half-maximal clustering during reversion.

### DNA-FISH

Cells were transfected with sgRNAs in a 384-well microplate. The next day, cells were illuminated with a 455-nm LED for 3 h as detailed above, followed by fixation with paraformaldehyde (10 min, RT). After three washes with PBS, cells were permeabilized with 0.5% Triton X-100 and 0.5% saponin in PBS (20 min, RT), washed twice with PBS, incubated with 0.1 N HCl (15 min, RT), and neutralized with 2X saline-sodium citrate (SSC) buffer (5 min, RT). Then, cells were pre-hybridized with 2X SSC, 50% formamide (overnight, 4°C). Fluorescently labeled BAC probes (Table S3) and commercial hybridization buffer were purchased from Empire Genomics (Buffalo, NY, USA). BAC probes were diluted in this buffer following the manufacturer’s instructions and added to the cells. Cells were then subjected to DNA denaturation (8 min, 85°C) followed immediately by hybridization (overnight, 37°C). The next day, hybridization mix was solubilized by adding 2X SSC (5 min, RT), cells were washed with 1X SSC (3 x 5 min, 42°C) and with 0.1X SSC (3 x 5 min, 42°C). Finally, cells were stained with 1 µg/mL DAPI (10 min, RT), washed and kept in PBS. The plate was subjected to high-throughput imaging or stored at 4°C for a few days before imaging.

### RNA-FISH

Cells were transfected with sgRNAs in a 384-well microplate. The next day cells were illuminated with a 455-nm LED for 4 h as detailed above, and fixed with paraformaldehyde (10 min, RT). After three washes with PBS, cells were permeabilized with 70% ethanol (2 h, RT), washed three times with PBS, and equilibrated with RNA-FISH wash buffer (2X SSC, 10% formamide) for 30 min at RT. Stellaris® RNA FISH probes for intronic *TERT* pre-mRNA were purchased as a pool of 30 oligonucleotides labeled with Atto565 from LGC Biosearch Technologies (Hoddesdon, UK) and are listed in Table S4. Hybridization was performed overnight at 37°C with 10 nM probe, 2X SSC, 10% formamide, 10% dextran sulfate. Cells were then washed with RNA-FISH wash buffer (30 min, 37°C + 5 min, RT), PBS (2 x 5 min, RT), and stained with 1 µg/mL DAPI (10 min, RT). The plate with cells kept in PBS was subjected to high-throughput imaging or stored at 4°C for a few days before imaging.

### RNA/DNA-FISH

Cells were subjected to RNA-FISH as described above and immediately subjected to high-throughput imaging (see below). Immediately after imaging, DNA-FISH was performed as described above, except for HCl treatment (7 min instead of 15 min) and pre-hybridization (1h, RT instead of overnight, 4°C). The plate was imaged again, acquiring images of the same fields imaged before.

### Image acquisition

For the analysis of CRY2 variants clustering and kinetic properties, cellular samples were imaged with a confocal laser scanning microscope LSM880 (Zeiss, Oberkochen, Germany), using a Plan-Apochromat 63x/1.4 Oil DIC M27 objective. mCherry was excited with a 594 nm HeNe laser, and CRY2 was activated with an Ar laser at 488 nm. Fluorescence filtered for the 598-696 nm range was detected with a GaAsP detector. Single-plane images of 512 x 512 pixels were acquired, with a pixel size of 80 nm. Signal was averaged 16 times with a final frame time of 5 s.

For other experiments, samples were imaged with a CV8000 high-throughput spinning disk confocal microscope (Yokogawa, Tokyo, Japan), using a 60x water objective (NA=1.2). DAPI, GFP/Fluorescein, mCherry/Atto565, and Cy5 were excited with 405 nm, 488 nm, 561 nm, and 640 nm lasers, respectively, and a 405/488/561/640 excitation dichroic mirror. Fluorescence was filtered through emission bandpass filters (445/45 nm, 525/50 nm, 600/37 nm, and 676/29 nm, respectively) and detected by two sCMOS cameras (2048 x 2048 pixels). *Z*-stacks spanning a range of 9 µm with 0.5 µm-intervals were acquired with a pixel size of 0.108 µm. Maximum *z*-projections, background correction, shading correction, and geometric corrections were performed on the fly by the proprietary Yokogawa software controlling the CV8000.

### Segmentation of nuclei and FISH spots

DNA-FISH and RNA-FISH images were directly analyzed as described below. RNA/DNA-FISH data required an additional initial image registration step to align the images of the same fields acquired sequentially on separate days. This RNA/DNA-FISH image alignment was performed as described previously (Almansour et al., 2024) with an algorithm based on Fourier phase correlation between the DAPI images corresponding to sequential RNA and DNA-FISH images to determine a spatial translation vector for alignment. The alignment was performed individually for each field. The image registration code is publicly available on GitHub (https://github.com/CBIIT/DNA_RNA_registration).

All RNA, DNA, and RNA/DNA-FISH data were analyzed with the open-source High-Throughput Image Processing Software (HiTIPS) (Keikhosravi et al., 2024) using the computational resources of the NIH HPC Biowulf cluster (https://hpc.nih.gov). Nuclei were segmented based on the DAPI channel using the deep learning model CellPose (Stringer et al., 2021) implemented for GPUs. FISH spots were segmented with the Gaussian Laplacian method as described in (Keikhosravi et al., 2024). Data from the same experiment was analyzed with the same segmentation parameters. The output of HiTIPS segmentation consisted of tables with single-nucleus and single-spot data, including size, shape, intensity, and spatial coordinates. Output data is publicly available on Figshare (see ‘Data and resource availability’).

### Analysis of FISH data

Output data from HiTIPS was analyzed with R (R version 4.4.1, https://cran.r-project.org). Scripts are publicly available on GitHub (see ‘Data and resource availability’). For quality control, nuclei smaller than 50 µm^2^ (HeLa) / 90 µm^2^ (U2OS) or with solidity < 0.9 were filtered out. For each nucleus, the number of FISH spots was calculated for each channel.

For DNA-FISH data, nuclei were also filtered out to only include cells with the correct number of FISH spots and according to the ploidy of each genome region (triploid for Subtel/*TERT* in HeLa, tetraploid for Subtel/*TERT* in U2OS, diploid for *IDR3/TCF3* in U2OS). All possible distances between DNA-FISH spots from different channels in the same cell were computed using the spot centroids *x,y* coordinates. Minimum distances below 2 µm were considered as the single-allele distances between the targeted DNA loci pairs. An arbitrary threshold of a distance of less than 0.27 µm was set to classify alleles as DNA contacts. This threshold reflects spatial resolution limitations since this is a typical DNA-FISH spot radius, and it is within the optical resolution limited by diffraction; similar thresholds have previously been used to measure chromatin interactions (Finn et al., 2019). Other thresholds generated different absolute values but did not significantly change results of experimental comparisons.

For RNA-FISH, the number of RNA spots per cell and the mean spot intensity were computed.

For RNA/DNA-FISH experiments, data was analyzed at the single-cell level. For each cell, the number of RNA spots, the sum of RNA spot intensities, the fraction of DNA distances < 0.27 µm, and the GFP nuclear intensity corresponding to dCas9-3XGFP-CRY2 levels were calculated. To facilitate the analysis and representation of data, cells were binned either to the fraction of close alleles per cell or the GFP intensity. Bins were set to maintain similar numbers of cells in each bin.

### Analysis of single-cell heterogeneity

Heterogeneity in the fraction of close alleles was analyzed at the single-cell level by computing for each experimental condition the fraction of cells with none, one, two or three alleles with distances < 0.27 µm, depending on the ploidy (diploid for *IDR3/TCF3* in U2OS, triploid for Subtel/*TERT* in HeLa). This observed distribution was compared with the expected distribution assuming that alleles are completely independent of each other. The expected fractions of close alleles *f*(*c*) were calculated for each condition according to the standard binomial distribution equation as follows:

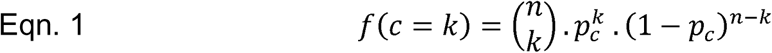

Where is *p_c_* is the frequency of close alleles in the population calculated for each condition from observed data, *n* is the ploidy of cells, and *k* is the number of close alleles to evaluate. From the above general binomial distribution formula, the expected fractions were calculated for diploid (Eqn. 2) and triploid (Eqn. 3) cell lines as follows:

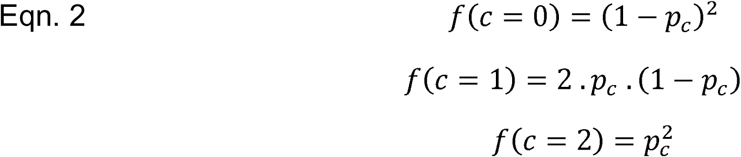

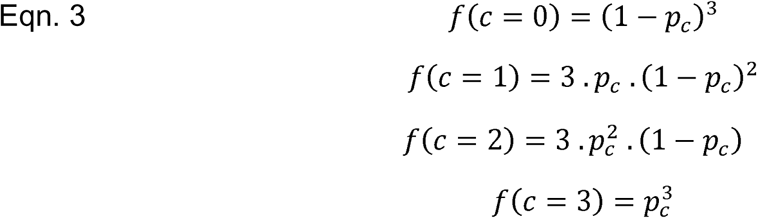

Scripts for these analyses are publicly available on GitHub (see ‘Data and resource availability’).

### Statistical analysis

Data was obtained from multiple independent experiments performed and measured on different days. Only relevant statistical comparisons are indicated in the figures with asterisks as * for p<0.05, ** for p<0.01, *** for p<0.001. Statistical analyses were done with scripts publicly available on GitHub (see ‘Data and resource availability’). For multiple comparisons of means involving two factors (for example, sgRNA and light treatments), a two-way ANOVA was performed followed by post-hoc Tukey tests. For comparing means of two conditions, t tests were performed. For comparing the observed distributions of cells with variable numbers of close alleles with the expected random distribution, a chi-square test was performed. When indicated, data from representative experiments are shown. In these cases, statistical comparisons were made considering the number of replicates as the number of alleles or cells measured. The Marascuilo’s procedure (Marascuilo, 1966) was performed for multiple comparisons of proportions (i.e. fractions of close alleles).

## ACKNOWLEDGMENTS

High-throughput imaging was performed in the NCI-CCR High-Throughput Imaging Facility. Confocal imaging was performed in the NCI-CCR Optical Microscopy Core supported by Dr. Tatiana Karpova and Dr. David Ball. High-throughput imaging data was processed using the computational resources of the NIH HPC Biowulf cluster (https://hpc.nih.gov). Cells were sorted in the NCI-CCR Flow Cytometry Core with assistance from Kathy McKinnon and Sophia Brown. We thank all members of the Misteli group for discussion. This research was supported by the Intramural Research Program of the National Institutes of Health (NIH), National Cancer Institute NCI, Center for Cancer Research through grant 1-ZIA-BC010309 to T.M. and grant 1-ZIC-BC-011567 to HiTIF. The contributions of the NIH authors were made as part of their official duties as NIH federal employees, are in compliance with agency policy requirements, and are considered Works of the United States Government. However, the findings and conclusions presented in this paper are those of the authors and do not necessarily reflect the views of the NIH or the U.S. Department of Health and Human Services.

## AUTHOR CONTRIBUTIONS

M.S. and T.M. conceptualized the project, designed experiments, interpreted results, and wrote the manuscript. T.M. obtained funding and supervised the project. M.S. performed the experiments and analyzed the data with scripts developed by A.K. and G.P.. All authors edited and approved the manuscript.

## DATA AND RESOURCE AVAILABILITY

Datasets and R scripts used to generate the results shown in this article are publicly available in Figshare (https://figshare.com/projects/mistelilab_optoloop/267052) and GitHub (https://github.com/CBIIT/mistelilab-optoloop), respectively.

## REFERENCES

1. Akdemir, K. C., Le, V. T., Chandran, S., Li, Y., Verhaak, R. G., Beroukhim, R., Campbell, P. J., Chin, L., Dixon, J. R., Futreal, P. A., et al. 2020. Disruption of chromatin folding domains by somatic genomic rearrangements in human cancer. Nat Genet, 52, 294–305.

2. Alexander, J. M., Guan, J., Li, B., Maliskova, L., Song, M., Shen, Y., Huang, B., Lomvardas, S. & Weiner, O. D. 2019. Live-cell imaging reveals enhancer-dependent Sox2 transcription in the absence of enhancer proximity. Elife, 8.

3. Almansour, F., Keikhosravi, A., Pegoraro, G. & Misteli, T. 2024. Allele-level visualization of transcription and chromatin by high-throughput imaging. Histochem Cell Biol, 162, 65–77.

4. Bartman, C. R., Hamagami, N., Keller, C. A., Giardine, B., Hardison, R. C., Blobel, G. A. & Raj, A. 2019. Transcriptional Burst Initiation and Polymerase Pause Release Are Key Control Points of Transcriptional Regulation. Mol Cell, 73, 519–532 e4.

5. Bartman, C. R., Hsu, S. C., Hsiung, C. C., Raj, A. & Blobel, G. A. 2016. Enhancer Regulation of Transcriptional Bursting Parameters Revealed by Forced Chromatin Looping. Mol Cell, 62, 237–247.

6. Benabdallah, N. S., Williamson, I., Illingworth, R. S., Kane, L., Boyle, S., Sengupta, D., Grimes, G. R., Therizols, P. & Bickmore, W. A. 2019. Decreased Enhancer-Promoter Proximity Accompanying Enhancer Activation. Mol Cell, 76, 473–484 e7.

7. Braselmann, E., Rathbun, C., Richards, E. M. & Palmer, A. E. 2020. Illuminating RNA Biology: Tools for Imaging RNA in Live Mammalian Cells. Cell Chem Biol, 27, 891–903.

8. Calderon, L., Weiss, F. D., Beagan, J. A., Oliveira, M. S., Georgieva, R., Wang, Y. F., Carroll, T. S., Dharmalingam, G., Gong, W., Tossell, K., et al. 2022. Cohesin-dependence of neuronal gene expression relates to chromatin loop length. Elife, 11.

9. Chakraborty, S., Kopitchinski, N., Zuo, Z., Eraso, A., Awasthi, P., Chari, R., Mitra, A., Tobias, I. C., Moorthy, S. D., Dale, R. K., et al. 2023. Enhancer-promoter interactions can bypass CTCF-mediated boundaries and contribute to phenotypic robustness. Nat Genet, 55, 280–290.

10. Chevalier, R., Murcia Pienkowski, V., Jullien, N., Caron, L., Pinard, P. V., Magdinier, F. & Robin, J. D. 2025. Telomere Position Effect-Over Long Distances Acts as a Genome-Wide Epigenetic Regulator Through a Common Alu Element. Aging Cell, 24, e70027.

11. Clow, P. A., Du, M., Jillette, N., Taghbalout, A., Zhu, J. J. & Cheng, A. W. 2022. CRISPR-mediated multiplexed live cell imaging of nonrepetitive genomic loci with one guide RNA per locus. Nat Commun, 13, 1871.

12. Conte, M., Esposito, A., Vercellone, F., Abraham, A. & Bianco, S. 2023. Unveiling the Machinery behind Chromosome Folding by Polymer Physics Modeling. Int J Mol Sci, 24.

13. Corin, A., Nora, E. P. & Ramani, V. 2025. Beyond genomic weaving: molecular roles for CTCF outside cohesin loop extrusion. Curr Opin Genet Dev, 90, 102298.

14. Deng, W., Lee, J., Wang, H., Miller, J., Reik, A., Gregory, P. D., Dean, A. & Blobel, G. A. 2012. Controlling long-range genomic interactions at a native locus by targeted tethering of a looping factor. Cell, 149, 1233–44.

15. Deng, W., Rupon, J. W., Krivega, I., Breda, L., Motta, I., Jahn, K. S., Reik, A., Gregory, P. D., Rivella, S., Dean, A., et al. 2014. Reactivation of developmentally silenced globin genes by forced chromatin looping. Cell, 158, 849–860.

16. Du, M., Zou, F., Li, Y., Yan, Y. & Bai, L. 2022. Chemically Induced Chromosomal Interaction (CICI) method to study chromosome dynamics and its biological roles. Nat Commun, 13, 757.

17. Duan, L., Hope, J., Ong, Q., Lou, H. Y., Kim, N., McCarthy, C., Acero, V., Lin, M. Z. & Cui, B. 2017. Understanding CRY2 interactions for optical control of intracellular signaling. Nat Commun, 8, 547.

18. Finn, E. H., Pegoraro, G., Brandao, H. B., Valton, A. L., Oomen, M. E., Dekker, J., Mirny, L. & Misteli, T. 2019. Extensive Heterogeneity and Intrinsic Variation in Spatial Genome Organization. Cell, 176, 1502–1515 e10.

19. Fulco, C. P., Nasser, J., Jones, T. R., Munson, G., Bergman, D. T., Subramanian, V., Grossman, S. R., Anyoha, R., Doughty, B. R., Patwardhan, T. A., et al. 2019. Activity-by-contact model of enhancer-promoter regulation from thousands of CRISPR perturbations. Nat Genet, 51, 1664–1669.

20. Grubert, F., Srivas, R., Spacek, D. V., Kasowski, M., Ruiz-Velasco, M., Sinnott-Armstrong, N., Greenside, P., Narasimha, A., Liu, Q., Geller, B., et al. 2020. Landscape of cohesin-mediated chromatin loops in the human genome. Nature, 583, 737–743.

21. Hernandez-Candia, C. N., Wysoczynski, C. L. & Tucker, C. L. 2019. Advances in optogenetic regulation of gene expression in mammalian cells using cryptochrome 2 (CRY2). Methods, 164-165, 81–90.

22. Hnisz, D., Weintraub, A. S., Day, D. S., Valton, A. L., Bak, R. O., Li, C. H., Goldmann, J., Lajoie, B. R., Fan, Z. P., Sigova, A. A., et al. 2016. Activation of proto-oncogenes by disruption of chromosome neighborhoods. Science, 351, 1454–1458.

23. Horsfield, J. A. 2023. Full circle: a brief history of cohesin and the regulation of gene expression. FEBS J, 290, 1670–1687.

24. Hsieh, T. S., Cattoglio, C., Slobodyanyuk, E., Hansen, A. S., Darzacq, X. & Tjian, R. 2022. Enhancer-promoter interactions and transcription are largely maintained upon acute loss of CTCF, cohesin, WAPL or YY1. Nat Genet, 54, 1919–1932.

25. Hyle, J., Zhang, Y., Wright, S., Xu, B., Shao, Y., Easton, J., Tian, L., Feng, R., Xu, P. & Li, C. 2019. Acute depletion of CTCF directly affects MYC regulation through loss of enhancer-promoter looping. Nucleic Acids Res, 47, 6699–6713.

26. Ing-Simmons, E., Vaid, R., Bing, X. Y., Levine, M., Mannervik, M. & Vaquerizas, J. M. 2021. Independence of chromatin conformation and gene regulation during Drosophila dorsoventral patterning. Nat Genet, 53, 487–499.

27. Jager, K., Mensch, J., Grimmig, M. E., Neuner, B., Gorzelniak, K., Turkmen, S., Demuth, I., Hartmann, A., Hartmann, C., Wittig, F., et al. 2022. A conserved long-distance telomeric silencing mechanism suppresses mTOR signaling in aging human fibroblasts. Sci Adv, 8, eabk2814.

28. Kane, L., Williamson, I., Flyamer, I. M., Kumar, Y., Hill, R. E., Lettice, L. A. & Bickmore, W. A. 2022. Cohesin is required for long-range enhancer action at the Shh locus. Nat Struct Mol Biol, 29, 891–897.

29. Keikhosravi, A., Almansour, F., Bohrer, C. H., Fursova, N. A., Guin, K., Sood, V., Misteli, T., Larson, D. R. & Pegoraro, G. 2024. High-throughput image processing software for the study of nuclear architecture and gene expression. Sci Rep, 14, 18426.

30. Kim, J. H., Rege, M., Valeri, J., Dunagin, M. C., Metzger, A., Titus, K. R., Gilgenast, T. G., Gong, W., Beagan, J. A., Raj, A., et al. 2019. LADL: light-activated dynamic looping for endogenous gene expression control. Nat Methods, 16, 633–639.

31. Kim, W., Ludlow, A. T., Min, J., Robin, J. D., Stadler, G., Mender, I., Lai, T. P., Zhang, N., Wright, W. E. & Shay, J. W. 2016. Regulation of the Human Telomerase Gene TERT by Telomere Position Effect-Over Long Distances (TPE-OLD): Implications for Aging and Cancer. PLoS Biol, 14, e2000016.

32. Kim, Y. J., Lee, M., Jr., Lee, Y. T., Jing, J., Sanders, J. T., Botten, G. A., He, L., Lyu, J., Zhang, Y., Mettlen, M., et al. 2023. Light-activated macromolecular phase separation modulates transcription by reconfiguring chromatin interactions. Sci Adv, 9, eadg1123.

33. Kragesteen, B. K., Spielmann, M., Paliou, C., Heinrich, V., Schopflin, R., Esposito, A., Annunziatella, C., Bianco, S., Chiariello, A. M., Jerkovic, I., et al. 2018. Dynamic 3D chromatin architecture contributes to enhancer specificity and limb morphogenesis. Nat Genet, 50, 1463–1473.

34. Krefting, J., Andrade-Navarro, M. A. & Ibn-Salem, J. 2018. Evolutionary stability of topologically associating domains is associated with conserved gene regulation. BMC Biol, 16, 87.

35. Levo, M., Raimundo, J., Bing, X. Y., Sisco, Z., Batut, P. J., Ryabichko, S., Gregor, T. & Levine, M. S. 2022. Transcriptional coupling of distant regulatory genes in living embryos. Nature, 605, 754–760.

36. Liu, J., Zhu, S., Hu, W., Zhao, X., Shan, Q., Peng, W. & Xue, H. H. 2023. CTCF mediates CD8+ effector differentiation through dynamic redistribution and genomic reorganization. J Exp Med, 220.

37. Lupianez, D. G., Kraft, K., Heinrich, V., Krawitz, P., Brancati, F., Klopocki, E., Horn, D., Kayserili, H., Opitz, J. M., Laxova, R., et al. 2015. Disruptions of topological chromatin domains cause pathogenic rewiring of gene-enhancer interactions. Cell, 161, 1012–1025.

38. Lyu, X. Y., Deng, Y., Huang, X. Y., Li, Z. Z., Fang, G. Q., Yang, D., Wang, F. L., Kang, W., Shen, E. Z. & Song, C. Q. 2022. CRISPR FISHer enables high-sensitivity imaging of nonrepetitive DNA in living cells through phase separation-mediated signal amplification. Cell Res, 32, 969–981.

39. Ma, H., Tu, L. C., Naseri, A., Chung, Y. C., Grunwald, D., Zhang, S. & Pederson, T. 2018. CRISPR-Sirius: RNA scaffolds for signal amplification in genome imaging. Nat Methods, 15, 928–931.

40. Ma, L., Guan, Z., Wang, Q., Yan, X., Wang, J., Wang, Z., Cao, J., Zhang, D., Gong, X. & Yin, P. 2020. Structural insights into the photoactivation of Arabidopsis CRY2. Nat Plants, 6, 1432–1438.

41. Mach, P., Kos, P. I., Zhan, Y., Cramard, J., Gaudin, S., Tunnermann, J., Marchi, E., Eglinger, J., Zuin, J., Kryzhanovska, M., et al. 2022. Cohesin and CTCF control the dynamics of chromosome folding. Nat Genet, 54, 1907–1918.

42. Marascuilo, L. A. 1966. Large-sample multiple comparisons. Psychol Bull, 65, 280–90.

43. Mas, P., Devlin, P. F., Panda, S. & Kay, S. A. 2000. Functional interaction of phytochrome B and cryptochrome 2. Nature, 408, 207–11.

44. Misteli, T. 2020. The Self-Organizing Genome: Principles of Genome Architecture and Function. Cell, 183, 28–45.

45. Morgan, S. L., Mariano, N. C., Bermudez, A., Arruda, N. L., Wu, F., Luo, Y., Shankar, G., Jia, L., Chen, H., Hu, J. F., et al. 2017. Manipulation of nuclear architecture through CRISPR-mediated chromosomal looping. Nat Commun, 8, 15993.

46. Narendra, V., Rocha, P. P., An, D., Raviram, R., Skok, J. A., Mazzoni, E. O. & Reinberg, D. 2015. CTCF establishes discrete functional chromatin domains at the Hox clusters during differentiation. Science, 347, 1017–21.

47. Nora, E. P., Goloborodko, A., Valton, A. L., Gibcus, J. H., Uebersohn, A., Abdennur, N., Dekker, J., Mirny, L. A. & Bruneau, B. G. 2017. Targeted Degradation of CTCF Decouples Local Insulation of Chromosome Domains from Genomic Compartmentalization. Cell, 169, 930–944 e22.

48. Oudelaar, A. M., Beagrie, R. A., Gosden, M., de Ornellas, S., Georgiades, E., Kerry, J., Hidalgo, D., Carrelha, J., Shivalingam, A., El-Sagheer, A. H., et al. 2020. Dynamics of the 4D genome during in vivo lineage specification and differentiation. Nat Commun, 11, 2722.

49. Ozkan-Dagliyan, I., Chiou, Y. Y., Ye, R., Hassan, B. H., Ozturk, N. & Sancar, A. 2013. Formation of Arabidopsis Cryptochrome 2 photobodies in mammalian nuclei: application as an optogenetic DNA damage checkpoint switch. J Biol Chem, 288, 23244–51.

50. Palayam, M., Ganapathy, J., Guercio, A. M., Tal, L., Deck, S. L. & Shabek, N. 2021. Structural insights into photoactivation of plant Cryptochrome-2. Commun Biol, 4, 28.

51. Park, E. J. & Kim, H. 2025. Live genome imaging by CRISPR engineering: progress and problems. Exp Mol Med, 57, 1392–1399.

52. Park, H., Kim, N. Y., Lee, S., Kim, N., Kim, J. & Heo, W. D. 2017. Optogenetic protein clustering through fluorescent protein tagging and extension of CRY2. Nat Commun, 8, 30.

53. Parmar, J. J., Woringer, M. & Zimmer, C. 2019. How the Genome Folds: The Biophysics of Four-Dimensional Chromatin Organization. Annu Rev Biophys, 48, 231–253.

54. Popay, T. M. & Dixon, J. R. 2022. Coming full circle: On the origin and evolution of the looping model for enhancer-promoter communication. J Biol Chem, 298, 102117.

55. Qin, G., Yang, J., Zhao, C., Ren, J. & Qu, X. 2022. Manipulating complex chromatin folding via CRISPR-guided bioorthogonal chemistry. Proc Natl Acad Sci U S A, 119, e2204725119.

56. Rao, S. S., Huntley, M. H., Durand, N. C., Stamenova, E. K., Bochkov, I. D., Robinson, J. T., Sanborn, A. L., Machol, I., Omer, A. D., Lander, E. S., et al. 2014. A 3D map of the human genome at kilobase resolution reveals principles of chromatin looping. Cell, 159, 1665–80.

57. Rao, S. S. P., Huang, S. C., Glenn St Hilaire, B., Engreitz, J. M., Perez, E. M., Kieffer-Kwon, K. R., Sanborn, A. L., Johnstone, S. E., Bascom, G. D., Bochkov, I. D., et al. 2017. Cohesin Loss Eliminates All Loop Domains. Cell, 171, 305–320 e24.

58. Robin, J. D., Ludlow, A. T., Batten, K., Magdinier, F., Stadler, G., Wagner, K. R., Shay, J. W. & Wright, W. E. 2014. Telomere position effect: regulation of gene expression with progressive telomere shortening over long distances. Genes Dev, 28, 2464–76.

59. Rodriguez, J., Ren, G., Day, C. R., Zhao, K., Chow, C. C. & Larson, D. R. 2019. Intrinsic Dynamics of a Human Gene Reveal the Basis of Expression Heterogeneity. Cell, 176, 213–226 e18.

60. Rowley, M. J. & Corces, V. G. 2018. Organizational principles of 3D genome architecture. Nat Rev Genet, 19, 789–800.

61. Schneider, N., Wieland, F. G., Kong, D., Fischer, A. A. M., Horner, M., Timmer, J., Ye, H. & Weber, W. 2021. Liquid-liquid phase separation of light-inducible transcription factors increases transcription activation in mammalian cells and mice. Sci Adv, 7.

62. Stringer, C., Wang, T., Michaelos, M. & Pachitariu, M. 2021. Cellpose: a generalist algorithm for cellular segmentation. Nat Methods, 18, 100–106.

63. Strom, A. R., Kim, Y., Zhao, H., Chang, Y. C., Orlovsky, N. D., Kosmrlj, A., Storm, C. & Brangwynne, C. P. 2024. Condensate interfacial forces reposition DNA loci and probe chromatin viscoelasticity. Cell, 187, 5282–5297 e20.

64. Taslimi, A., Vrana, J. D., Chen, D., Borinskaya, S., Mayer, B. J., Kennedy, M. J. & Tucker, C. L. 2014. An optimized optogenetic clustering tool for probing protein interaction and function. Nat Commun, 5, 4925.

65. Uyehara, C. M. & Apostolou, E. 2023. 3D enhancer-promoter interactions and multi-connected hubs: Organizational principles and functional roles. Cell Rep, 42, 112068.

66. Wang, H., Xu, X., Nguyen, C. M., Liu, Y., Gao, Y., Lin, X., Daley, T., Kipniss, N. H., La Russa, M. & Qi, L. S. 2018. CRISPR-Mediated Programmable 3D Genome Positioning and Nuclear Organization. Cell, 175, 1405–1417 e14.

67. Wang, J., Yu, H., Ma, Q., Zeng, P., Wu, D., Hou, Y., Liu, X., Jia, L., Sun, J., Chen, Y., et al. 2021. Phase separation of OCT4 controls TAD reorganization to promote cell fate transitions. Cell Stem Cell, 28, 1868–1883 e11.

68. Winick-Ng, W., Kukalev, A., Harabula, I., Zea-Redondo, L., Szabo, D., Meijer, M., Serebreni, L., Zhang, Y., Bianco, S., Chiariello, A. M., et al. 2021. Cell-type specialization is encoded by specific chromatin topologies. Nature, 599, 684–691.

69. Yang, J. H. & Hansen, A. S. 2024. Enhancer selectivity in space and time: from enhancer-promoter interactions to promoter activation. Nat Rev Mol Cell Biol, 25, 574–591.

70. Yang, L. Z., Min, Y. H., Liu, Y. X., Gao, B. Q., Liu, X. Q., Huang, Y., Wang, H., Yang, L., Liu, Z. J. & Chen, L. L. 2024. CRISPR-array-mediated imaging of non-repetitive and multiplex genomic loci in living cells. Nat Methods, 21, 1646–1657.

71. Zhu, Y., Balaji, A., Han, M., Andronov, L., Roy, A. R., Wei, Z., Chen, C., Miles, L., Cai, S., Gu, Z., et al. 2025. High-resolution dynamic imaging of chromatin DNA communication using Oligo-LiveFISH. Cell, 188, 3310–3328 e27.

