## Supplementary Materials for "OptoLoop: An optogenetic tool to probe the functional role of genome organization"

### SUPPLEMENTARY FIGURES

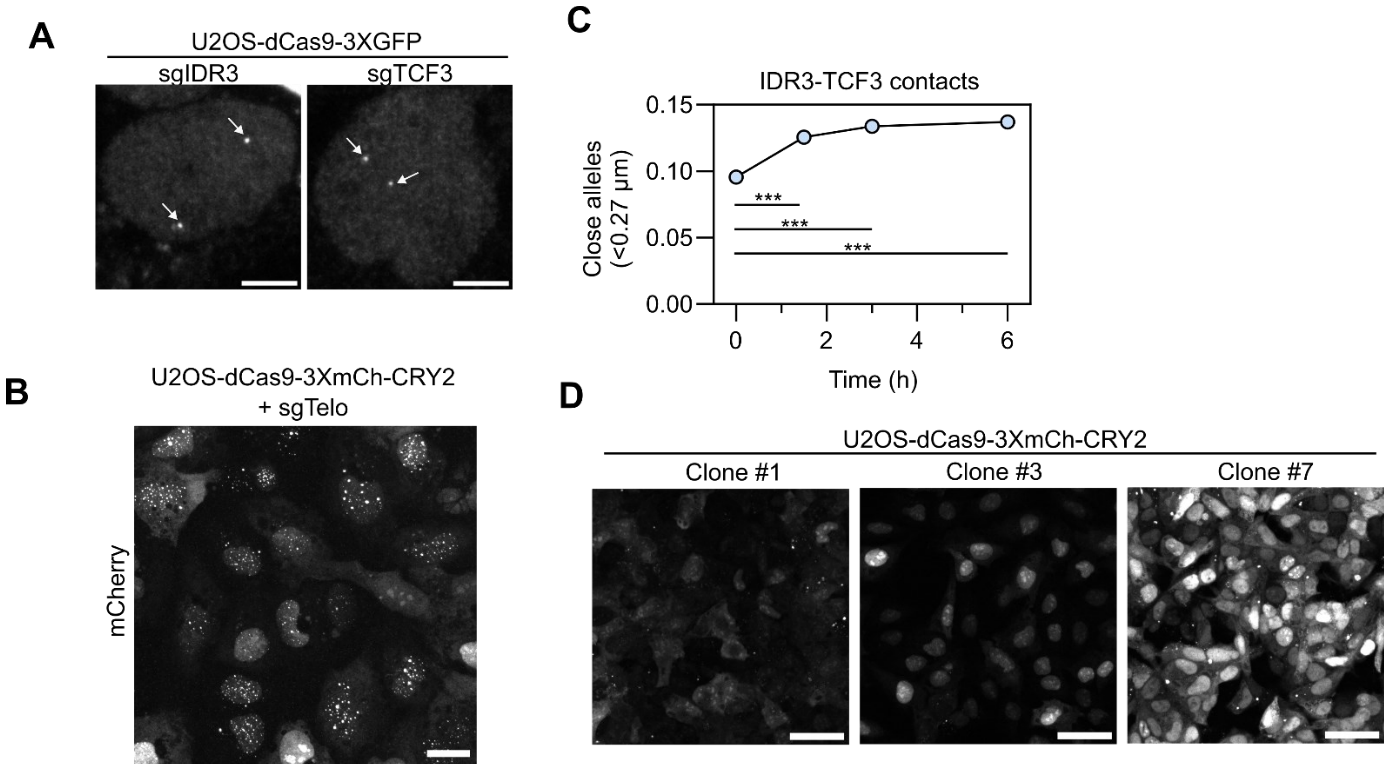


**Fig. S1. Setting up OptoLoop in U2OS cells. A)** Images of U2OS cells stably expressing dCas9-3XGFP and transfected with sgIDR3 and sgTCF3. White arrows indicate IDR3 and TCF3 labelled loci. Scale bar: 5 μm. **B)** Images of U2OS stably expressing dCas9-3XmCh-CRY2 (clone 3) and transfected with sgTelo. Scale bar: 20 μm. **C)** Fraction of alleles with IDR3-TCF3 distance < 0.27 μm measured from DNA-FISH images of U2OS cells stably expressing dCas9-3XmCh-CRY2 (clone 1), transfected with sgIDR3 and sgTCF3 and illuminated for variable times (1 s pulses every 10 s). Data from one experiment with typically 12000-15000 alleles analyzed per sample. **D)** Images of U2OS clones stably expressing different levels of dCas9-3XmCh-CRY2. All images are shown with the same intensity scale for comparison. Scale bar: 50 μm. Asterisks indicate significantly different comparisons (* for p<0.05, ** for p<0.01, *** for p<0.001).

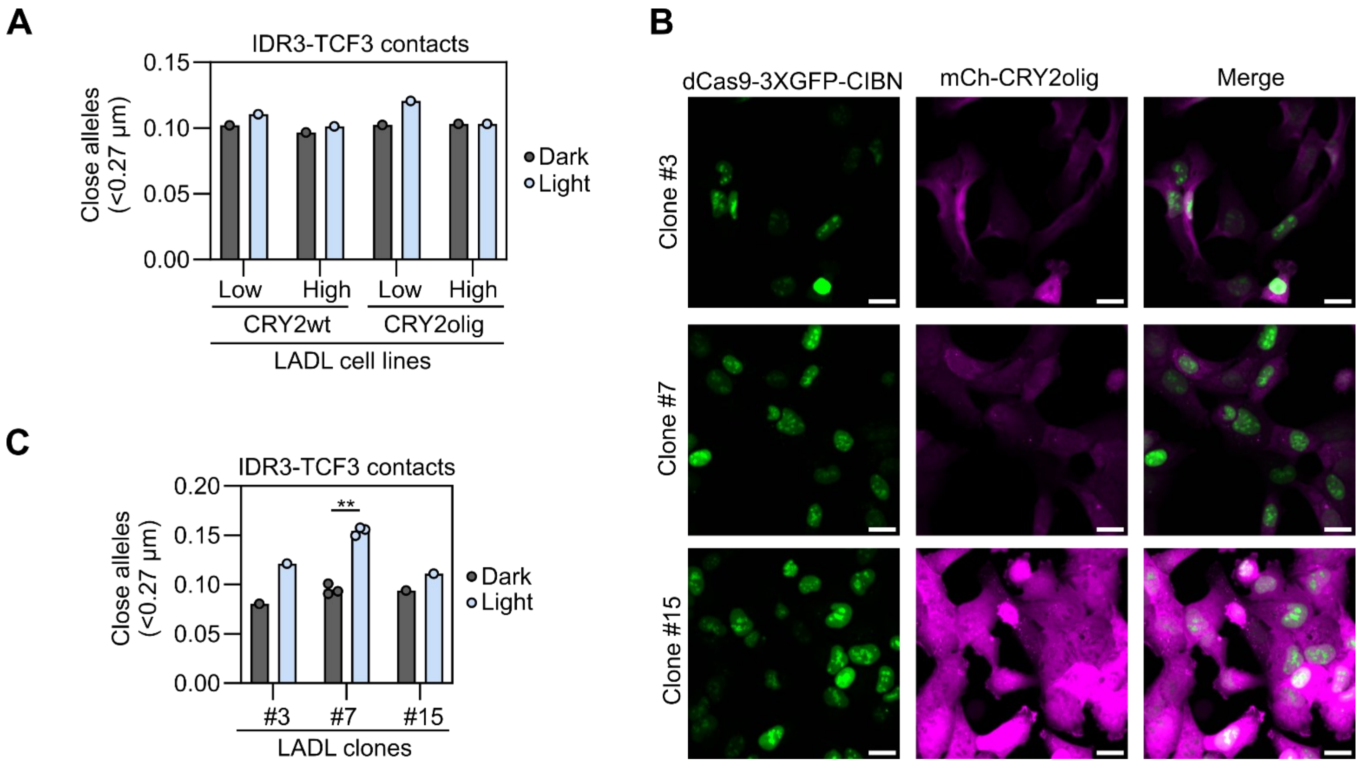


**Fig. S2. Setting up LADL in U2OS cells. A)** Fraction of alleles with IDR3-TCF3 distance < 0.27 µm measured from DNA-FISH images of polyclonal U2OS cell lines stably expressing dCas9-3XGFP-CRY2 and low/high expression levels of mCherry fused to CRY2wt or CRY2olig, transfected with sgIDR3 and sgTCF3, and kept under dark or illuminated with blue light for 3 h (1 s pulses every 10 s). Data from one experiment with typically 3000-5000 alleles analyzed per sample. **B)** Images of U2OS clones stably expressing different levels of dCas9-3XGFP-CIBN and mCherry-CRY2olig. All images are shown with the same intensity scale for comparison. Scale bar: 20 µm. **C)** Fraction of alleles with IDR3-TCF3 distance < 0.27 µm measured from DNA-FISH images of U2OS clones stably expressing dCas9-3XGFP-CIBN and variable levels of mCherry-CRY2olig, transfected with sgIDR3 and sgTCF3, and kept under dark or illuminated with blue light for 3 h (1 s pulses every 10 s). Each dot represents the fraction of typically 3500-7000 alleles analyzed per experiment. Bars represent the means of independent experiments. Asterisks indicate significantly different comparisons (* for p<0.05, ** for p<0.01, *** for p<0.001).

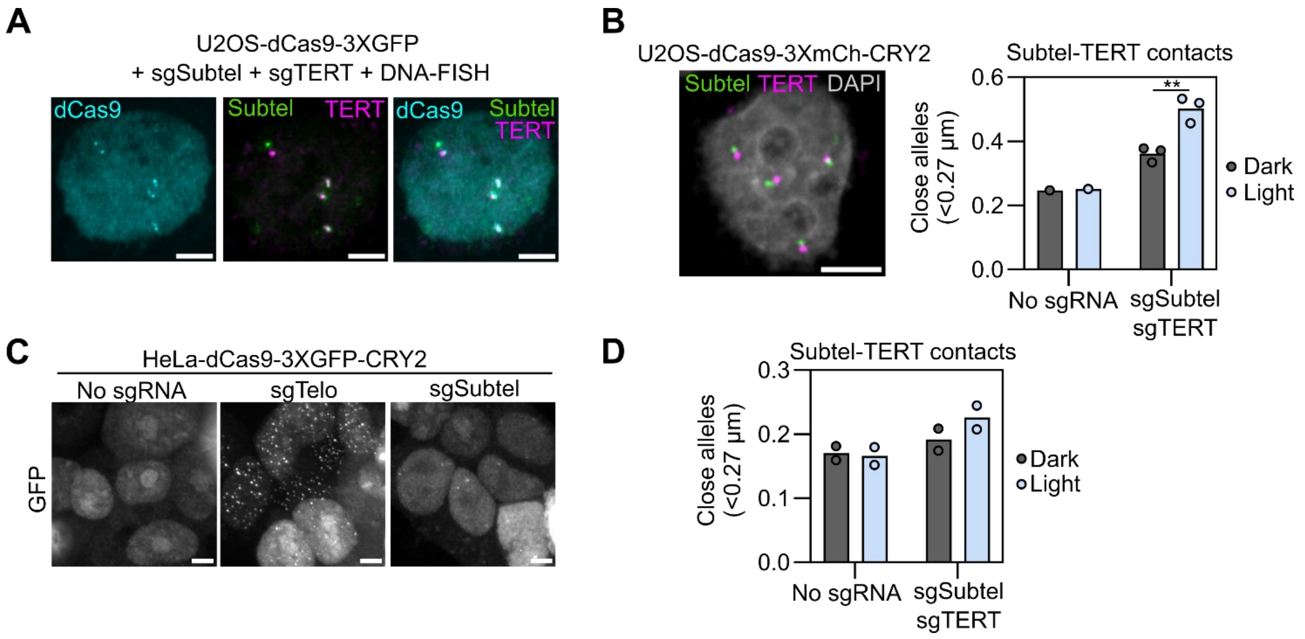


**Fig. S3. Manipulation of Subtel-TERT contacts in U2OS and HeLa cells. A)** Sequential images of a U2OS cell stably expressing dCas9-3XGFP and transfected with sgSubtel and sgTERT, first imaged for GFP, and then imaged after DNA FISH with Subtel and TERT BAC probes. Scale bar: 5 µm. **B)** Left panel: Representative image of DNA FISH with Subtel and TERT BAC probes in U2OS. Scale bar: 5 µm. Right panel: Fraction of alleles with Subtel-TERT distance <0.27 µm measured from DNA-FISH images of U2OS cells stably expressing dCas9-3XmCherry-CRY2 (clone 3), transfected with none or sgSubtel and sgTERT sgRNAs, and kept under dark or illuminated with blue light for 3 h (1 s pulses every 10 s). Each dot represents the fraction of typically 5000-10000 alleles analyzed per experiment. Bars represent the means of independent experiments. Asterisks indicate significantly different comparisons (* for p<0.05, ** for p<0.01, *** for p<0.001). **C)** Images of HeLa cells stably expressing dCas9-3XGFP-CRY2 and transfected with the indicated sgRNAs. Scale bar: 5 µm. **D)** Fraction of alleles with Subtel-TERT distance <0.27 µm measured from DNA-FISH images of HeLa cells stably expressing dCas9-3XGFP-CRY2, transfected with none or sgSubtel and sgTERT sgRNAs, and kept under dark or illuminated with blue light for 3 h (1 s pulses every 10 s). Each dot represents the fraction of typically 6000-20000 alleles analyzed per experiment. Bars represent the means of two experiments.

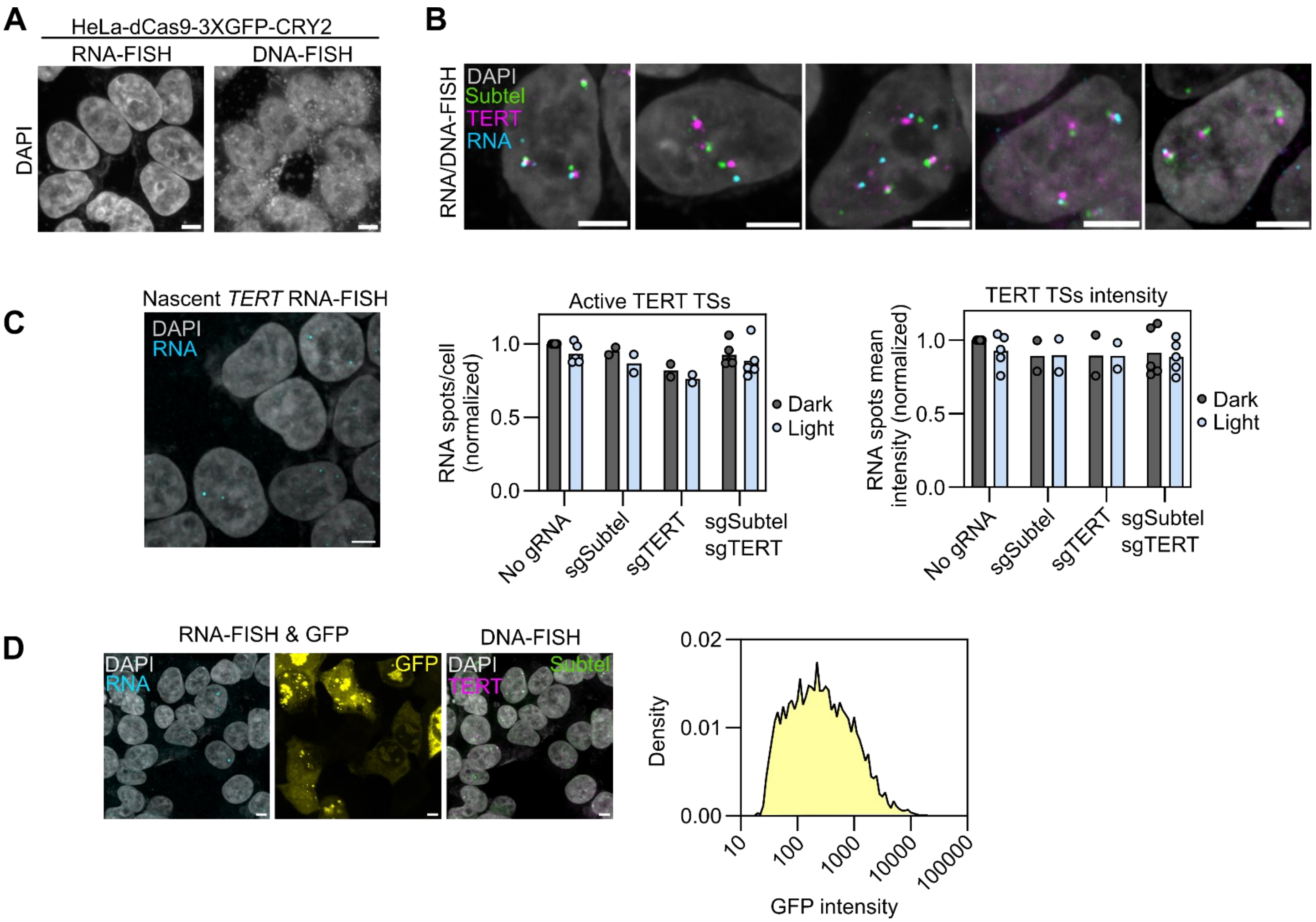


**Fig. S4. Analysis of DNA looping and RNA transcription. A)** Images of HeLa cells after RNA-FISH and after sequential DNA-FISH. DAPI staining is shown. Scale bar: 5 µm. **B)** Examples of RNA/DNA-FISH images of HeLa cells with TERT pre-mRNA probes and Subtel and TERT BAC probes. Scale bar: 5 µm.**C)** Left panel: Image of nascent RNA-FISH with probes against TERT intron in HeLa cells. Scale bar: 5 µm. Middle and right panels: Number of active TERT transcription sites per cell and their mean intensity, respectively, measured from RNA-FISH images from HeLa cells stably expressing dCas9-3XGFP-CRY2, transfected with the indicated combinations of sgSubtel and sgTERT, and illuminated with blue light for 4 h (1 s pulses every 10 s). Values are represented as relative to control (no sgRNAs, no light). Each dot represents the mean of typically 3000-12000 cells analyzed per experiment. Bars represent the means of independent experiments. **D)** Left panels: Images of sequential RNA/DNA FISH of HeLa cells stably expressing dCas9-3XGFP-CRY2 with TERT pre-mRNA probes and Subtel and TERT BAC probes. GFP-channel was acquired simultaneously with RNA FISH imaging, before DNA FISH. Right panel: GFP nuclear intensity histogram. Scale bar: 10 µm.

### SUPPLEMENTARY TABLES

**Table S1. List of plasmids.**

| Plasmid | Source | Reference |
|---|---|---|
| pCMV-CRY2olig-mCherry | Addgene #60032 | (Taslimi et al., 2014) |
| pCMV-mCherry-CRY2clust | Addgene #105624 | (Park et al., 2017) |
| pCMV-CRY2high-mCherry | Addgene #104063 | (Duan et al., 2017) |
| pCMV-CRY2hiclu-mCherry | Site-directed mutagenesis | This paper |
| pHAGE-dCas9-3XGFP | Addgene #64107 | (Ma et al., 2015) |
| pHAGE-dCas9-3XGFP-CRY2 | High-fidelity assembly cloning | This paper |
| pHAGE-dCas9-3XmCherry | Addgene #64108 | (Ma et al., 2015) |
| pHAGE- dCas9-mCherry-CRY2 | PCR/restriction cloning | This paper |
| pHAGE-dCas9-3XmCherry-CRY2 | High-fidelity assembly cloning | This paper |
| pHR-mCherry-CRY2wt | Addgene #101221 | (Shin et al., 2017) |
| pHR-mCherry-CRY2olig | Addgene #101222 | (Shin et al., 2017) |
| pEF1a-dCas9-CIBN | Addgene #127664 | (Kim et al., 2019) |
| pHAGE-dCas9-3XGFP-CIBN | PCR/restriction cloning | This paper |
| psPAX2 | Addgene #12260 | Gift from Didier Trono (unpublished) |
| pMD2.G | Addgene #12259 | Gift from Didier Trono (unpublished) |

**Table S2. List of single-guide RNAs.** Ordered as synthetic RNAs from Synthego.

| Name | Spacer sequence | # repeats | Location |
|---|---|---|---|
| sgIDR3 | UGAUGAGCAGAUGUAGGAGG | 34 | chr19: 380,831-382,654 |
| sgTCF3 | AAGGGGACAGCAGAGCUCAC | 35 | chr19: 1,627,754-1,628,973 |
| sgTelo | UUAGGGUUAGGGUUAGGGUU | ~$10^2$-$10^3$ | Telomeres |
| sgSubtel1 | CGAUGAGAGGCCGGGUGGUC | 32 | chr5: 83,314-85,466 |
| sgSubtel2 | CGGGAGGCGGAGCUCACA | 97 | chr5: 258,622-262,350 |
| sgTERT1 | GUGUGCUGGGGGCUCACUGG | 58 | chr5: 1,069,995-1,073,282 |
| sgTERT2 | UGUGUCUGUAGAGGAGGAGC | 35 | chr5: 1,326,332-1,328,969 |

**Table S3. List of BACs for DNA FISH.** Ordered labeled and ready to use from Empire Genomics.

| Name | Clone | Dye | Location (hg19) |
|---|---|---|---|
| IDR3 | RP11-575H1 | Cy5-dUTP | chr19: 258,325-423,094 |
| TCF3 | RP11-317H11 | Fluorescein-dUTP | chr19: 1,380,124-1,544,933 |
| Subtel | RP11-44H14 | Fluorescein-dUTP | chr5:93,896-269,773 |
| TERT | RP11-117B23 | Cy5-dUTP | chr5:1,207,044-1,369,139 |

**Table S4. List of nascent RNA FISH probes for TERT.** Ordered as a pool and labeled with Atto565 from LGC Biosearch Tecnhologies.

| **Probe sequence** | **Target** |
|---|---|
| ccgaccccggggaggcccac | Intron 1 |
| cacgtgacgatggagacagg | Intron 2 |
| tcgacgtcctgagcgaaaag | Intron 2 |
| gaacctcgtaagtttatgca | Intron 2 |
| aaaccgcgtgtccatcaaaa | Intron 2 |
| attataggtaacctgcaggc | Intron 2 |
| cacccttgaaattgcgaaga | Intron 2 |
| tgtgctggagaacagtctta | Intron 2 |
| ccctcttcatacctaaagat | Intron 2 |
| ggttcctcaacatcaaatcc | Intron 2 |
| ggtgaaatcgggacttcttc | Intron 2 |
| tcttacatgtcttgggagtt | Intron 2 |
| catgacgcttatctgactcg | Intron 2 |
| cgaaggaagctggagcacaa | Intron 2 |
| agtcctgatcagagaactca | Intron 2 |
| ctcaagtgagcaaacgccaa | Intron 2 |
| aaaagataggcttggggacc | Intron 2 |
| aatggaacggagaggtgacc | Intron 2 |
| caagctgggagaggagtatt | Intron 2 |
| agagaaccctttctgggatg | Intron 2 |
| gcagcttgggagaaaacagg | Intron 2 |
| tgcccagaataaggtgacaa | Intron 2 |
| tggacgtcaatccatgtgag | Intron 2 |
| ctaagacccaagagggaagt | Intron 2 |
| gcacttagagggaaaggcat | Intron 2 |
| tcaaaacacagggtgcaggt | Intron 2 |
| gtggctgatgttgagattac | Intron 2 |
| caggtaaagctggggtttac | Intron 2 |
| tgtgcatcataagcagaggt | Intron 2 |
| cactcactggctaaggaacg | Intron 2 |
| caggtttgcgcgatttcaaa | Intron 2 |

**SUPPLEMENTARY REFERENCES**

Duan, L., Hope, J., Ong, Q., Lou, H. Y., Kim, N., McCarthy, C., Acero, V., Lin, M. Z. & Cui, B. 2017. Understanding CRY2 interactions for optical control of intracellular signaling. *Nat Commun,* 8**,** 547.

Kim, J. H., Rege, M., Valeri, J., Dunagin, M. C., Metzger, A., Titus, K. R., Gilgenast, T. G., Gong, W., Beagan, J. A., Raj, A., et al. 2019. LADL: light-activated dynamic looping for endogenous gene expression control. *Nat Methods,* 16**,** 633-639.

Ma, H., Naseri, A., Reyes-Gutierrez, P., Wolfe, S. A., Zhang, S. & Pederson, T. 2015. Multicolor CRISPR labeling of chromosomal loci in human cells. *Proc Natl Acad Sci U S A,* 112**,** 3002-7.

Park, H., Kim, N. Y., Lee, S., Kim, N., Kim, J. & Heo, W. D. 2017. Optogenetic protein clustering through fluorescent protein tagging and extension of CRY2. *Nat Commun,* 8**,** 30.

Shin, Y., Berry, J., Pannucci, N., Haataja, M. P., Toettcher, J. E. & Brangwynne, C. P. 2017. Spatiotemporal Control of Intracellular Phase Transitions Using Light-Activated optoDroplets. *Cell,* 168**,** 159-171 e14.

Taslimi, A., Vrana, J. D., Chen, D., Borinskaya, S., Mayer, B. J., Kennedy, M. J. & Tucker, C. L. 2014. An optimized optogenetic clustering tool for probing protein interaction and function. *Nat Commun,* 5**,** 4925.
